# Probing the conserved roles of Cut in the development and function of optically different insect compound eyes

**DOI:** 10.1101/2022.10.14.512307

**Authors:** Shubham Rathore, Michael Meece, Mark Charlton-Perkins, Tiffany A. Cook, Elke K Buschbeck

## Abstract

Astonishing functional diversity exists among arthropod eyes, yet eye development relies on deeply conserved genes. This phenomenon is best understood for early events, whereas fewer investigations have focused on the influence of later transcriptional regulators on diverse eye organizations and the contribution of critical support cells, such as Semper cells (SCs). As SCs in *Drosophila melanogaster* secrete the lens and function as glia, they are critical components of ommatidia. Here, we perform RNAi-based knockdowns of the transcription factor *cut* (CUX in vertebrates), a marker of SCs, the function of which has remained untested in these cell types. To probe for the conserved roles of *cut*, we investigate two optically different compound eyes: the apposition optics of *D. melanogaster* and the superposition optics of the diving beetle *Thermonectus marmoratus*. In both cases, we find that multiple aspects of ocular formation are disrupted, including lens facet organization and optics as well as photoreceptor morphogenesis. Together, our findings highlight a generalized role for SCs in arthropod eye form and function and define Cut as a central player in mediating these functions.

## Introduction

Compound eyes, a prominent eye type in arthropods, are known for their typically highly organized multifaceted convex corneal lenses and underlying photoreceptor (PR) clusters that are components of precisely organized visual units (ommatidia). Despite sharing similar features, compound eye types show remarkable diversity, often related to the animal’s ecology (Land and Nilsson, 2012; Cronin *et al*., 2014; Meece *et al*., 2021; Nilsson, 2021). Optically, compound eyes can be divided into two general types: apposition and superposition (Nilsson, 1983, 1989; Meyer-Rochow, 2015). In the more ancestral apposition eye, each lens only serves its own underlying PRs, as exemplified in *Drosophila melanogaster* (Fig. 1A). Note that *D. melanogaster* eyes are actually neural-superposition eyes, which refers to a neural (rather than optical) organization that allows pooling from neighboring units (Nilsson, 1989, 2021; Agi *et al*., 2014). Consequently, from a strictly optical perspective, apposition organization allows each lens to project a tiny inverted image onto the underlying PR array. Therefore, neighboring units function relatively independently. The second compound eye type is the more derived superposition eye as exemplified by the Sunburst Diving Beetle *Thermonectus marmoratus* (Fig. 1B). In this eye type many lenses are precisely organized to allow the synergistic projection of images onto the underlying PR array (Nilsson, 1989), which increases light sensitivity (Warrant, 1999). To facilitate optical pooling, this organization requires a clear zone between the optical components and a more distally placed rhabdom layer. Because large parts of the eye must work together, even small changes in the arrangement can lead to optical deficiencies. Despite these functional and structural differences, the cellular organization of ommatidia is relatively conserved, with few differences among arthropods (Paulus, 1979).

**Fig. 1.**
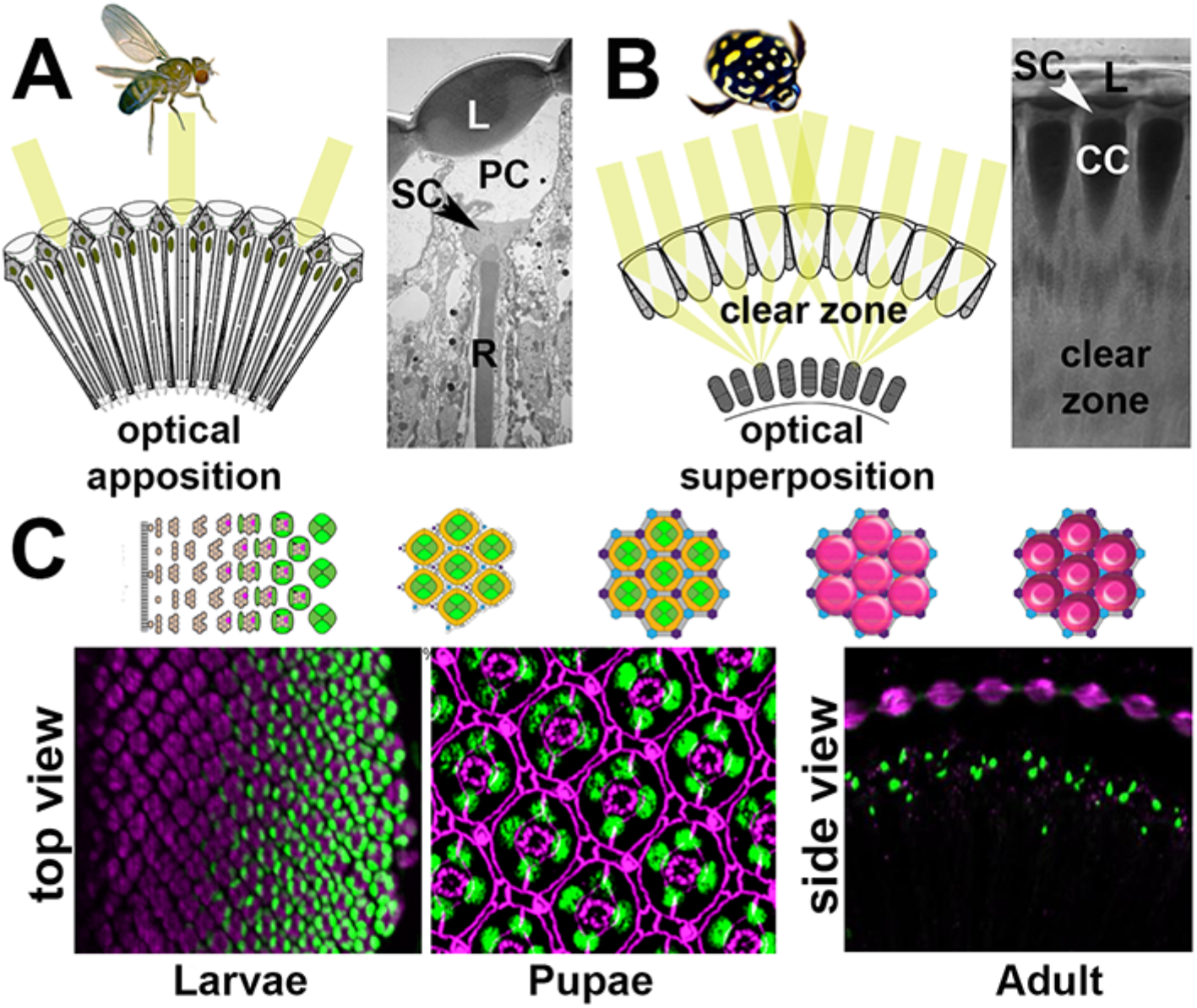
Compound eye types and development. A. *D. melanogaster* has a neural-superposition eye, the optics of which follows typical apposition organization, with individual lenses (L) that each project a tiny image fragment onto the tips of underlying photoreceptor rhabdomeres (R). Underneath the lens, there is a pseudocone (PC) and four Semper cell (SC) bodies. B. *T. marmoratus* has an optical superposition eye, in which sets of lenses synergistically project image points onto corresponding underlying closed rhabdoms. In this organization, the SC bodies are located in close proximity to the lens and above the photoreceptor cell bodies. The optics require the presence of pronounced crystalline cones (CCs) and a clear zone. Compound eye development is best understood in *D. melanogaster*, in which specific cell types are sequentially recruited from a precursor epithelium. Top: Diagram of cell fate specification and differentiation in the *Drosophila* compound eye. Bottom: The four SCs within each ommatidium show Cut immunoreactivity (green) in the larval, pupal, and adult stages. For better orientation, counterstained tissue is illustrated in magenta: ELAV in larvae, E-cadherin in pupae, and drosocrystallin (lenses) in adults.

Building on genetic studies in *D. melanogaster*, the development of insect compound eyes has been shown to share similar patterns, irrespective of eye type (Friedrich, 2003). For example, PRs develop first from a precursor epithelium, followed by accessory cell development and ultimately lens secretion, pigment cell differentiation, and photoreceptor morphogenesis (Buschbeck and Friedrich, 2008). This process is best understood in the compound eyes of *D. melanogaster*. Due to their precise crystalline organization, and ease of genetic manipulation, the development of these eyes has become a key model for studying organogenesis and tissue patterning. Forming from an undifferentiated neuro-epithelial imaginal disc of a late third instar larva, an eye develops that has ∼800 ommatidia, each containing eight PRs, four Semper (cone) cells (SCs), and two primary pigment cells (PPCs), all surrounded by pigmented epithelium-like tissue (composed of higher-order pigment cells) that optically isolates individual ommatidia (Fig. 1C)(Charlton-Perkins and Cook, 2010).

At the molecular level, several genes, including *pax6*, have been identified as being part of an ancestrally conserved gene network (the retinal determination network, RDGN) that underlies early eye development (Gehring, 2001; Kumar and Moses, 2001; Mishra and Sprecher, 2020). The genes in the RDGN are important for specifying the early developing cells of ommatidia and are known to be well conserved among some species (Gehring, 2001; Kumar, 2001; Hsiung and Moses, 2002; Mishra and Sprecher, 2020). A few studies have highlighted the nuances of how conserved expression patterns affect early eye development in different species (Yang *et al*., 2009*a*,*b*). However, relatively little is known about how conserved genes that can influence later eye development processes, such as the delineation of ommatidia as discrete units, act in different compound eyes. One gene of interest is the homeodomain transcription factor *cut*, which in *D. melanogaster* eyes is known for its specific expression in SCs.

SCs are recruited immediately after PRs through a combination of extrinsic signaling factors and intrinsic transcription factors (Charlton-Perkins *et al*., 2021). Two such factors that are essential to the coordination of these events are Pax2 and Prospero, which cooperatively specify SCs and later independently control PR structure and function. SCs are in close proximity to PRs and are a key component of ommatidia (Fig. 1B) with multiple roles, including recruiting PPCs, patterning the interommatidial cells, secreting the lens and pseudocone, and serving structural and functional support roles for retinal photoreceptors (Waddington and Perry, 1960; Cagan and Ready, 1989; Querenet *et al*., 2015; Charlton-Perkins *et al*., 2017, 2021; Stahl *et al*., 2017*b*). Their organization into a quartet is an evolutionarily conserved feature of arthropod compound eyes (Richter, 2002; Schwentner *et al*., 2018; Scholtz *et al*., 2019). In *D. melanogaster*, the transcriptional regulation of SC fate determination is relatively well understood (Cagan and Ready, 1989; Kumar, 2012; Morrison *et al*., 2018; Charlton-Perkins *et al*., 2021) and their differentiation follows closely that of PRs. As they secrete part of the corneal lens and the pseudocone during the second half of development, SCs are important candidates for influencing the optics of developing eyes. Evidence for their glial nature stems from physiology, which demonstrates ionic and metabolic support for PRs, and from their molecular profile (Cagan and Ready, 1989; Charlton-Perkins *et al*., 2017). SCs also express important conserved transcription factors, such as Pax2, which is upstream of Cut (Fu and Noll, 1997; Charlton-Perkins *et al*., 2021).

Cut is a particularly interesting transcription factor because it is a known marker of SCs within *D. melanogaster* ommatidia (Fu and Noll, 1997; Charlton-Perkins *et al*., 2011). It has been established that the SC-specific expression of Cut lasts throughout the life of *D. melanogaster* (Fig. 1C)(Charlton-Perkins *et al*., 2011). In SCs, Cut has been shown to partner with other transcription factors like Lozenge and Groucho to repress the expression of developmentally relevant genes like *deadpan* (Canon and Banerjee, 2003). Outside eyes, Cut is generally known for its developmental role in cell growth regulation and cell type differentiation (Nepveu, 2001). In *D. melanogaster*, it controls early cell fate decisions in most embryonic tissues, including the nervous system (Blochlinger *et al*., 1993). In external sensory organs, *cut* specifically belongs to a network of genes that are essential for the accurate maturation of specific sensory neurons and perineurial glia (Blochlinger *et al*., 1991; Bauke *et al*., 2015; Corty *et al*., 2016). Additionally, Cut prevents chordotonal cell fates in these external sensory organs (Blochlinger *et al*., 1991) and is required for the accurate morphogenesis of other sensory organs such as mechanosensory bristles and auditory organs in *D. melanogaster* (Hardiman et al., 2002; Ebacher et al., 2007) and *Tribolium castaneum* (Klann et al., 2021). Despite the known important developmental roles in the *D. melanogaster* nervous system, the function of Cut within SCs remains largely unexplored.

To investigate the potential contribution of Cut in SCs to the development of two very different compound eye types (apposition optics in *D. melanogaster* and superposition optics in *T. marmoratus*), we first established that Cut expression is conserved in SCs. We then conducted a comparative loss-of-function study and found remarkably similar knockdown phenotypes; this is consistent with the idea that a versatile shared developmental pathway underlies functionally different eye types.

## Results

### Cut expression is conserved within the four SCs of *D. melanogaster* and *T. marmoratus* compound eyes and can be successfully knocked down in both species

In *D. melanogaster*, Cut is expressed in the 4 SCs of each ommatidium and the interommatidial mechanosensory bristle (Fig 2A & Supp Fig. 1A) (Blochlinger *et al*., 1990). Our immunohistochemical investigation using the *D. melanogaster* antibody found it also expressed the 4 SCs of *T*.*marmoratus*, suggesting a high level of protein conservation between the two species (Fig 2B). N-cadherin and DAPI (for *D. melanogaster*) or actin and DAPI (for *T. marmoratus*) were used to confirm the position of the SC cell bodies distal to the developing PRs and rhabdoms. These similarities led us to hypothesize a conserved functional role for this transcription factor between these two eye types. To test for such conserved roles, we used a loss-of-function approach, applying a Semper cell-restricted knockdown in *D. melanogaster* (Charlton-Perkins et al., 2017) and a generalized dsRNAi injection in late stage *T. marmoratus* larvae (Rathore *et al*., 2020).

**Fig. 2.**
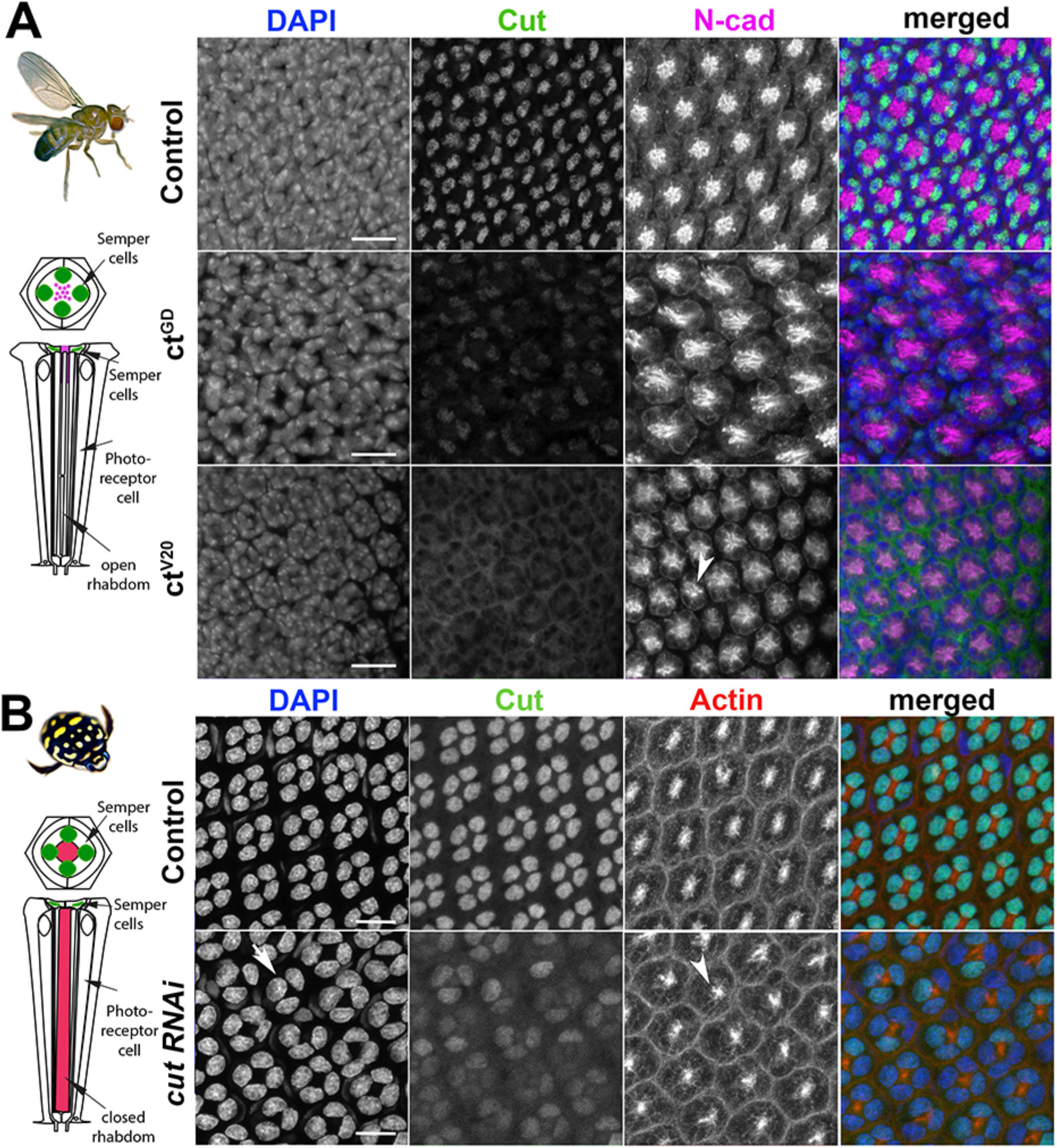
Cut expression and RNAi-driven knockdown in *D. melanogaster* and *T. marmoratus* compound eyes. A. In *D. melanogaster*, a quartet of Cut-positive Semper cells (green) are situated within the distal-most portion of developing ommatidia (at ∼37% pupal development). DAPI and N-cadherin counterstaining are used to identify the correct layer within the eye. As illustrated in representative images, efficient *cut* knockdown is achieved by two different SC-directed RNAis, with overlapping phenotypes consisting of an irregular ommatidial array. Incidences of laterally displaced rhabdoms are indicated via N-cadherin staining (arrowhead). B. In *T. marmoratus*, at a comparable developmental stage, four Cut-positive SCs are similarly organized near the distal margin of each ommatidium. At this stage, the closed rhabdom (red; confirmed by actin staining) still resides in close proximity to the SCs. The nuclear localization of Cut is confirmed by complete overlap with DAPI. *cut*RNAi treated individuals show a strong but incomplete reduction of Cut, with irregularities in the ommatidial array. In some instances at this level, only a triad of Cut-positive nuclei are visible (arrow), and rhabdoms appear to be laterally displaced (arrowhead). Scale bars = 10 μm.

To confirm the SC-restricted knockdown in *D. melanogaster*, we analyzed whole mounts of 37% developed pupal retina (which is prior to lens formation) in two different *cut*RNAi lines (*ct*^*GD*^ and *ct*^V20^) and compared them to a control line (*gfp-*RNAi). We found that SC-directed knockdowns reduced Cut expression either partially (*ct*^GD^; n = 14) or to an undetectable level (*ct*^V20^; n = 14, Fig. 2A) when compared to controls (n = 15). This difference could be due to the driver being heterozygous in *ct*^GD^ flies and homozygous in *ct*^V20^ flies, or to differences in knock-down efficiencies between these two lines. To verify that *cut* knockdown was restricted to SCs, we also imaged Cut-expressing interommatidial bristle cells located proximal to the SCs. As expected, Cut expression was unaffected in these cells in all three fly lines (Supp Fig. 1A).

To ensure a comparable stage and preparation for *T. marmoratus*, we injected *cut* dsRNA in late third instar larvae, and stained for Cut in early developing whole pupal retina. Consistent with previous findings (Rathore *et al*., 2020), we found knockdowns in *T. marmoratus* to be highly successful, with 10 of 12 injected individuals showing reduced Cut expression when compared to controls (n = 8). However, complete *cut* knockdown was rare, likely because of *cut*’s vital role in the development of multiple organ systems in insects (Bauke *et al*., 2015; Corty *et al*., 2016; Klann *et al*., 2021). Accordingly, ∼53% of the *cut* dsRNA injected individuals died, as opposed to only 27% of the controls. Although the SCs of control individuals strongly expressed Cut, the *cut* dsRNA-injected individuals exhibited SCs with varying, but greatly reduced Cut expression (Fig. 2B).

### Cut in SCs regulates precise ommatidial patterning during the development of different compound eye types

To investigate whether *cut* knockdowns affect the ommatidial organization during development, we examined pupal retinae for patterning defects. In *D. melanogaster*, N-cadherin allowed visualization of the position of the apical membranes of SCs and their relative position to PRs (Hayashi and Carthew, 2004). For both knockdown lines, we found visible organizational changes in these cell types as well as irregularities in the placement of ommatidia (Fig. 2A and Supp Fig. 1B). Typically, at ∼40% after puparium formation (APF), SCs are expected to be fully differentiated and organized in a characteristic tetrad, with the inner junctions forming an “H”-like pattern (see (Fu and Noll, 1997; Charlton-Perkins *et al*., 2021) and Control in Supp Fig. 1B). In contrast, the rhabdomeres in PRs are located within the center of each ommatidium and are expected to elongate toward the basement membrane (Longley and Ready, 1995). In both of *ct* knockdown lines, the typical “H” shape (indicative of a tetrad of SCs) failed to develop. Instead, an irregular pattern emerged with several instances of triads (Supp Fig. 1B). In parallel, the *cut*RNAi of the developing *T. marmoratus* pupal retina also showed irregularities and instances of SC triads (Fig. 2B). Although not verifiable with the N-cadherin antibody due to a lack of cross-reactivity, this phenomenon was clearly apparent with DAPI. The presence of triads in both species suggests that only three of the four SCs in these ommatidia developed properly, with one of the cells likely either absent or radically displaced.

Additionally, for *D. melanogaster*, control PRs had (as expected) centrally located rhabdomeres, whereas both *ct* knockdown lines showed displaced rhabdomere organization. Deficiencies included the dislocation of rhabdomeres from the central position of the ommatidium (Fig. 2A). In parallel, based on actin counterstaining of the actin-rich rhabdom structures, several of the fused rhabdoms in the *T. marmoratus cut*RNAi individuals also lost precise alignment at the center of the ommatidia relative to controls (Fig. 2B). Taken together, these observations show that *cut* knockdown in compound eyes results in complimentary developmental organizational deficiencies in both SCs and PRs. To investigate the functional implications of these developmental defects in adult compound eyes, we focused on the three known major functions of SCs: lens formation, PR and rhabdom morphogenesis, and PR function support.

### Cut is required by SCs for proper lens formation and function

Lens secretion begins at ∼50% pupal development of *D. melanogaster* retina, and the role of SCs in this process has been well documented (Cagan and Ready, 1989; Charlton-Perkins and Cook, 2010). Considering our findings that Cut disrupts accurate SC patterning, we next asked if these SC defects could lead to structural and functional defects in adult compound eye corneal lenses. To address this question, we first assessed the outer lens surfaces at an ultrastructural level. In both *D. melanogaster* (Fig. 3A–F) and *T. marmoratus* (Fig. 3G–K), knockdown individuals showed visible defects in lens morphology, shape, and organization when compared to controls.

**Fig. 3.**
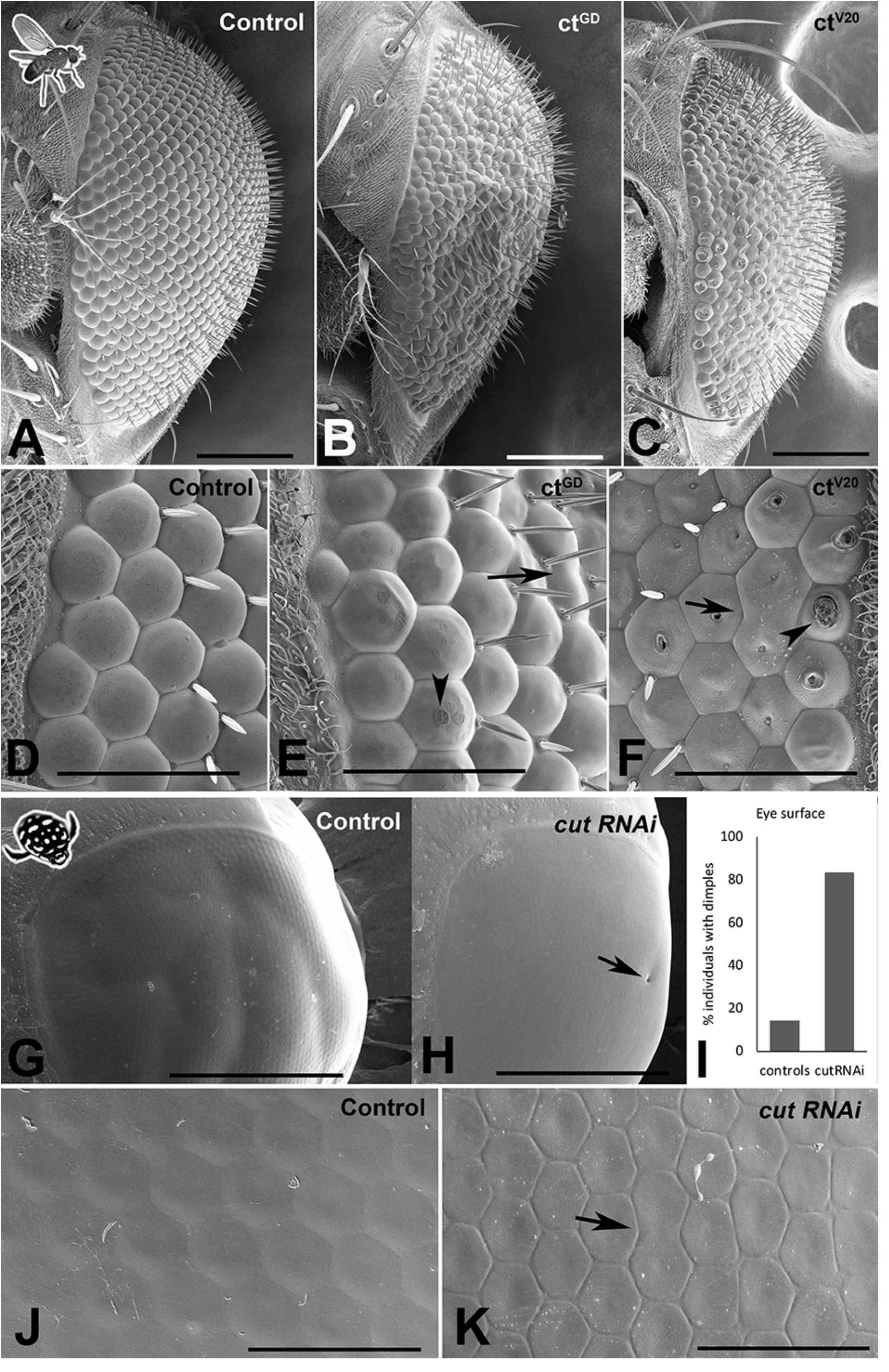
*Cut* knockdown affects lens organization in insect compound eyes. A–F. Scanning electron micrographs of adult *D. melanogaster* compound eyes. Overview of a control individual (A) illustrates a typical completely regular ommatidium array, whereas *ct*^GD^ (B) and *ct*^V20^ (C) exhibit major irregularities in ommatidial placement and lens formation. The latter is illustrated in a magnified view of the anterior region of the compound eye. In control individuals (D), lenses appear precisely shaped with properly formed lens surfaces. In *ct*^GD^ (E) and *ct*^V20^ (F), irregularities exist in ommatidium separation, with some neighboring units fused (arrows). In some instances, the lens surface exhibits deformities typical of the blueberry phenotype (arrowheads), which are particularly pronounced in the *ct*^V20^ line. G–K. Scanning electron micrographs of adult *T. marmoratus* compound eyes. Overview of a control beetle (G) shows an intact eye with a smooth surface. In *cut*RNAi individuals (H), surface dimples (arrow) are more common than in controls (I). A high-resolution image of the anterior region of the compound eye illustrates precise placement and smooth transitions between neighboring ommatidia (J), whereas *cut*RNAi injected individuals show irregularities in ommatidium size and more delineated borders (K). Additionally, some neighboring units are fused (arrow). Scale bars = 100 μm (A–C), 40 μm (D–F), 500 μm (G,H), and 50 μm (J,K).

In *D. melanogaster*, in contrast to the precisely organized lenses of controls (n = 9; Fig. 3A,D), both *cut* knockdown lines (*ct*^GD^: n = 10, *ct*^V20^: n = 14) showed major defects, including a rough eye appearance and variation among lenses (Fig. 3B,C). In both lines, we observed instances of fusion of neighboring units and flatter lenses, some of which had visible indentations on the outer lens surface (Fig. 3E,F), a phenotype previously referred to as “blueberry phenotype” (Higashijima *et al*., 1992). The latter phenomenon was particularly prominent in the *ct*^V20^ line. The position of the indentations were often asymmetric, possibly relating to differential contributions by different SCs.

Adult *T. marmoratus* compound eyes tended to have a smooth surface, which appeared similar in control and *cut*RNAi individuals at low resolution (Fig. 3G,H). However, the RNAi individuals (n = 6) showed an increase in dimple-like developmental defects on the outer eye surface (∼80% of individuals) when compared to controls (∼20%; n = 7; Fig. 3I). Higher resolution images revealed that unlike the highly regular and seamlessly connected lens arrays of controls (Fig. 3J), *cut*RNAi individuals generally had irregular lens shapes and pronounced ommatidial boundaries.

To assess lenses from a functional perspective, we isolated lens arrays and mounted them onto a droplet of insect ringer (hanging drop method). This approach allowed examination of the back surfaces of the lenses as well as the visualization of the corresponding images formed by the isolated lens arrays. In both species, control individuals (n = 6 for *D. melanogaster* and 10 for *T. marmoratus*) exhibited smooth lens surfaces (Fig. 4A,D), and the images produced by the lens arrays were regularly spaced with uniform image magnification (Fig. 4A’,D’). In contrast, in *D. melanogaster ct* knockdown lines, the aforementioned lens deficits were also visible from the back surface (n = 6 for each line; Fig. 4B,C). Notably, in *ct*^V20^ individuals, some lenses appeared necrotic (Iyer *et al*., 2016), while others showed small indentations. Optically, both lines showed instances where neighboring lenses formed images with irregular placement, focused at different planes, and exhibited variable image magnification (Fig. 4B’,C’). For the *ct*^V20^ line, we also noted that some lenses with holes that entirely failed to form images.

**Fig. 4.**
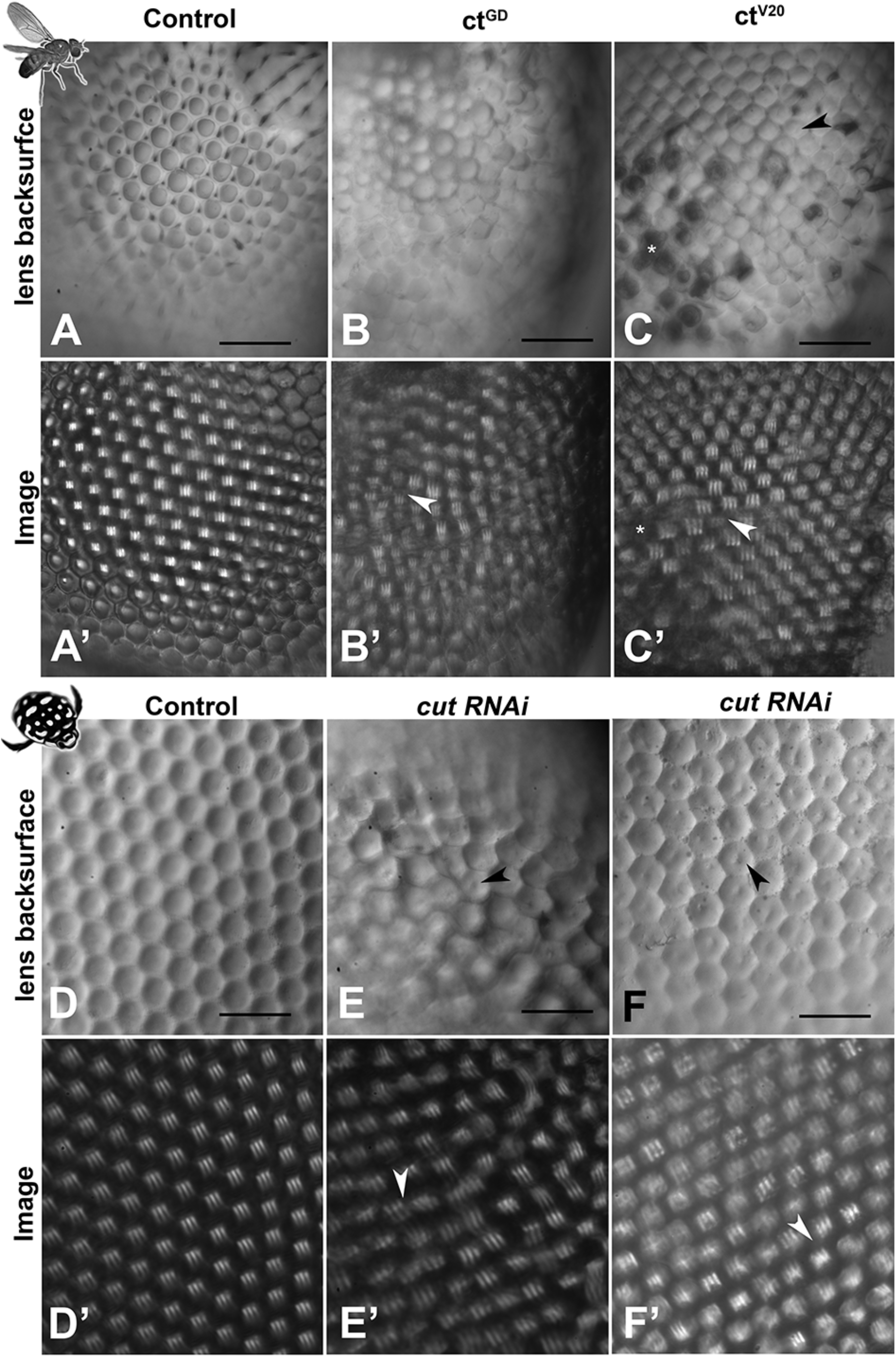
Morphological lens defects lead to profound optical deficits. Isolated lens arrays were used to visualize the back surfaces of lenses (A–F) and images of an object with three stripes were then produced by these lens arrays (A’–F’). In *D. melanogaster*, control lenses have smooth and accurately formed back surfaces (A) that lead to regularly spaced and equally sized images with a uniform focal plane across the lens array (A’). In contrast, the lenses of the *cut* knockdown lines show visible defects in morphology (B,C) and optics, with images that vary in placement, image magnification (arrowheads), focal plane, and blurriness (B’,C’). For *ct*^V20^, several lenses show dimple-like indentations on the back surfaces (arrowhead) and other lenses appear dark and necrotic (C). Such necrotic lenses (exemplified by the cluster marked with *) lead to gaps in the resulting image array (* in C’). In *T. marmoratus*, a similar pattern is observed, with controls having smooth and even lens back surfaces (D) that result in pristine regular image arrays (D’). In contrast, *cut*RNAi individuals exhibit lens irregularities (E) that lead to irregularities in the corresponding lens array (E’), including greatly displaced images (arrowhead). Lens back surfaces frequently show dimple-like lens indentations (arrowheads in E,F), which are also present in individuals with fewer irregularities in lens placement (F). Even in this morphologically less severe phenotype, major deficiencies in the lens array optics occur, resulting in many blurry images and some differently sized images (arrowhead) (F’). Scale bars = 50 μm.

Mirroring the findings from *D. melanogaster, cut* dsRNA injected *T. marmoratus* (n = 10) exhibited irregular lens arrays with centrally located indentations on the back surfaces (Fig. 4E,F). Many of the lens back surfaces also appeared flatter. Accordingly, the images produced by the lens arrays differed in placement, quality, and image magnification (Fig. 4E’,F’). Taken together, these results suggest that Cut is generally required by SCs for accurate lens formation with proper optics in both compound eye types.

### Cut is required by SCs for accurate PR placement and rhabdom morphogenesis

SCs are located in close proximity to PRs, and in *D. melanogaster*, they are known to provide glial support (ionic, metabolic, and structural) to PRs (Charlton-Perkins *et al*., 2017). The same study also demonstrated that SC-specific *pax2*RNAi results in defective rhabdom morphogenesis. Considering that Cut functions downstream of Pax2 and based on our observed displacements of the developing rhabdoms, *cut* knockdown could also lead to misformed adult rhabdoms.

It is important to note that *D. melanogaster* has open rhabdoms (each consisting of eight rhabdomeres) that extend distally close to the SC bodies (Fig. 5A,B), whereas in *T. marmoratus*, closed rhabdoms are separated from the SCs via a clear zone (Fig. 5E,F). As illustrated by phalloidin staining, despite these structural differences, we found similar knockdown phenotypes in the two species.

**Fig. 5.**
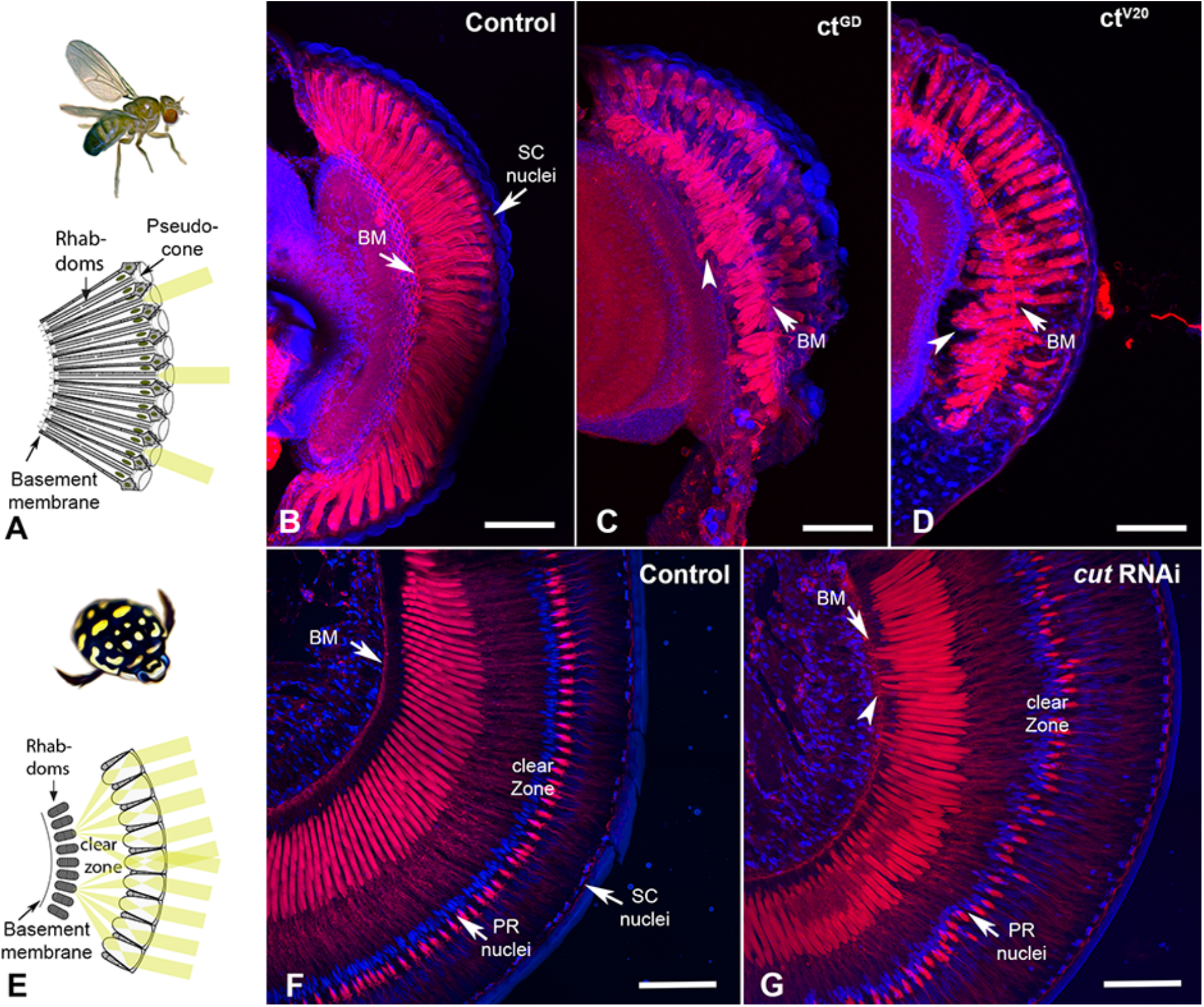
*Cut* knockdown leads to rhabdom misplacement in both eye types. A. In *D. melanogaster*, rhabdoms typically extend along the majority of the ommatidia, from close to the pseudocone to the basement membrane. B. Rhabdoms, visualized with phalloidin, appear well developed and regular in control individuals. In *ct*^GD^ (C) and *ct*^V20^ (D) individuals, rhabdoms appear truncated and frequently misplaced, with many extending well below the basement membrane (arrowheads). E. In *T. marmoratus*, rhabdoms are situated much deeper in the eye to make room for a clear zone, which is necessary to allow many lenses to contribute to the image formed at the distal end of the PRs. F. Phalloidin staining in control individuals illustrates precisely aligned rhabdoms that extend from below the clear zone to well above the basement membrane (BM). PR nuclei are aligned precisely along a concentric circle between the rhabdoms and lenses. G. In *cut*RNAi individuals, PR placement is less regular, at the levels of both PR nuclei and rhabdoms. As in *D. melanogaster*, rhabdoms are displaced toward the basement membrane and occasionally traverse it (arrowhead). Scale bars = 50 μm (B–D) and 100 μm (F,G).

In *D. melanogaster*, compared to the pristinely organized rhabdoms of control individuals that traverse the entire length of each ommatidium (n = 10; Fig. 5B), the rhabdoms for both *ct*^GD^ (n = 7) and *ct*^V20^ (n = 6) knockdown individuals were truncated. Generally, they either failed to reach the basement membrane or were present deep in the eye, either traversing the basement membrane or are seen proximally to it.

Similarly, in *T. marmoratus*, controls (n = 7) showed precisely placed PRs with a well-defined clear zone (Fig. 5F). However, the retina of *cut*RNAi individuals (n = 7) showed irregular PR placement (including misplaced nuclei), with some PRs extending to and even through the basement membrane (Fig. 5G). Additionally, the clear zone in these individuals was less well defined.

At an ultrastructural level, *D. melanogaster* controls (based on two TEM preparations) exhibited the classical trapezoid organization (Longley and Ready, 1995) with seven visible open rhabdomeres (Fig. 6A,B). In contrast, individuals from both the *ct*^GD^ (n = 3) and *ct*^V20^ (n = 2) lines exhibited parallel deficiencies, such as irregularly spaced ommatidia (Fig. 6C,F), a reduced number of rhabdomeres (Fig. 6C,H), split rhabdomeres (Fig. 6D,E,G), and fused rhabdomeres (Fig. 6H). For *T. marmoratus*, controls showed regularly organized closed rhabdoms, whereas rhabdom abnormalities were observed in *cut*RNAi individuals (Fig. 6I,J). The retina of *cut*RNAi individuals showed irregularly placed rhabdoms, with some units being on a different plane within the same array (Fig. 6K,N). Additionally, the rhabdoms were frequently incomplete and off center (Fig. 6M,O) with instances of split and broken manifestations (Fig. 6L,P) and signs of degeneration (Fig. 6P). Overall, these results suggest that Cut functions in SCs to direct the proper development of PRs in both compound eye types.

**Fig. 6.**
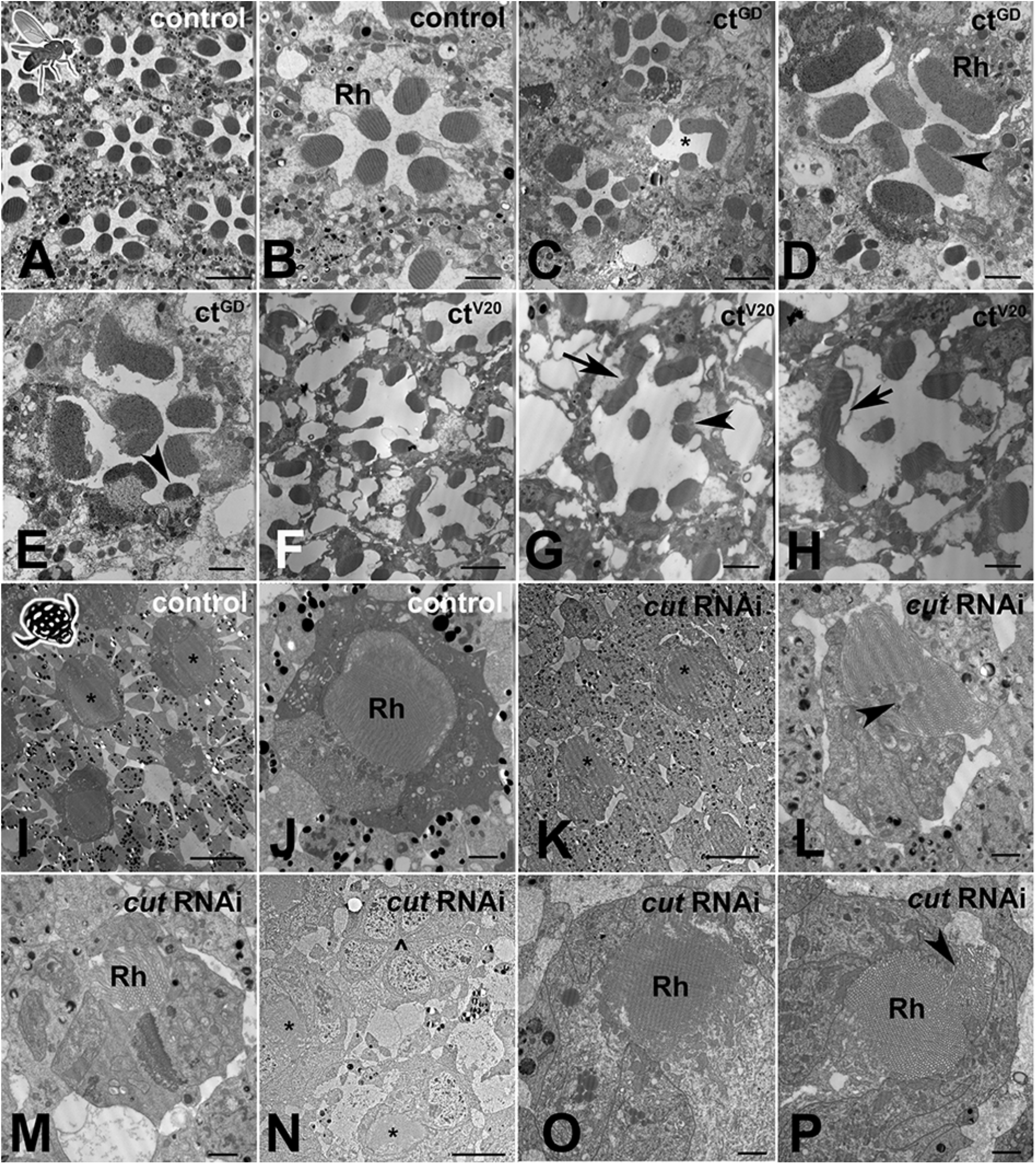
*Cut* knockdown leads to ultrastructural defects of rhabdoms in both eye types. A. As illustrated by a control individual, *D. melanogaster* has an open rhabdom that, at any cross-sectional plane, is formed by seven rhabdomeres. B. At higher magnification, it is apparent that the smaller central rhabdomere extends into the center of an extracellular lumen, which is bordered by larger and approximately evenly sized outer rhabdomeres. C. Overview of a *ct*^GD^ knockdown individual illustrates ommatidial displacements (with a compromised interommatidial space) and deformed or missing rhabdomeres (exemplified by the unit marked with *). D. Several units characterized by relatively extended or even split rhabdomeres (arrowhead). E. Other units showing unusually small rhabdomeres (arrowhead). F. Overview of a *ct*^V20^ knockdown individual illustrates ommatidia with relatively sparse rhabdomeres, large extracellular spaces between rhabdomeres, and sparse and degenerate interommatidial tissue. Nonetheless, *ct*^V20^ individuals also show laterally extended rhabdoms (G, arrow), split rhabdomeres (G, arrowhead), and possibly fused rhabdomeres (H, arrow). I. As illustrated by a control individual, the superposition eyes of *T. marmoratus* are characterized by closed rhabdoms (two units with similar rhabdom diameters in close proximity are marked with *). J. The rhabdom is positioned centrally within a healthy ommatidium. K. In *cut*RNAi individuals, neighboring units (marked with *) show relatively different rhabdom organization. L. An unusually shaped rhabdom with central deficiencies. M. A laterally displaced and strongly degenerate rhabdom. N. Overview of several ommatidia in a different individual shows the complete absence of a rhabdom (^) next to two neighboring semi-intact rhabdoms (*). O. A laterally degenerate rhabdom. P. A rhabdom with a displaced portion (arrowhead). Scale bars = 5 μm (A,C,F), 2 μm (B,D,E,G,H,J,L,M,O,P), and 10 μm (I,K,N); Rh = rhabdomere (B,D) or rhabdom (J,M,O,P).

### Cut in SCs does not majorly affect PR function in either species

To understand whether morphological defects related to *cut* knockdowns also affected function, we used electroretinograms (ERGs), an extracellular recording technique that measures the response of the PR array to light stimuli (Belusic, 2011; Riazuddin *et al*., 2012; Charlton-Perkins *et al*., 2017).

In *D. melanogaster*, we first assessed the dynamic ranges of PR responses to increasing light intensities (Fig. 7A) and found relatively normal responses in both knockdown lines (Fig. 7B). Fig. 7A illustrates the response curves for individuals from both control and *cut*RNAi lines to 490nm light stimuli of an intermediate light intensity (7.2 **×** 10^11^ photons/cm^2^/s). As expected from a normal fly ERG, control PR responses were flanked by on and off transients. Although flies from both *cut*RNAi lines had slightly reduced PR responses, there was no statistical difference at two different light intensities (Fig. 7C), which may, in part, be due to the high variability in all groups, which may be related to slight differences in eye color. In *T. marmoratus*, all controls had normal PR responses, but dramatic differences existed in *cut*RNAi individuals, which were separated into three categories (Fig. 7D): a) individuals with normally shaped responses at all light intensities, b) individuals with normally shaped responses at low light intensities but inverted responses at higher light intensities, and c) individuals with reversed-polarity responses at all light intensities. To keep the quantitative analysis comparable to controls, only individuals with electronegative responses were incorporated into the statistical analysis. Fig. 7E illustrates such normal and reversed potential responses. In some cases, in a train of pulses, only the first response was inverted (Fig. 7H). As shown in Fig. 7F, even the normal, electronegative responses of *cu*tRNAi individuals tended to be lower than those of controls (n = 15), with a significant difference at a light intensity of 10^13^ photons (n = 7; p = 0.0015, Wilcoxon’s test) but not at 4 **×** 10^13^ (n = 4; p = 0.08, Wilcoxon’s test) (Fig. 7G). Note that statistical power was lost at higher light intensities due to smaller sample sizes resulting from an increased occurrences of reversed polarity potentials (5.00 **×** 10^11^, n = 8; 1.90 **×** 10^12^, n = 8; 5.00 **×** 10^12^, n = 8; 1.00 **×** 10^13^, n = 7; 2.20 **×** 10^13^, n = 5; 5.50 **×** 10^13^, n = 5; 1.14 **×** 10^14^, n = 5; 2.25 **×** 10^14^, n = 4; 3.90 **×** 10^14^, n = 5; and 1.07 **×** 10^14^, n = 4).

**Fig. 7.**
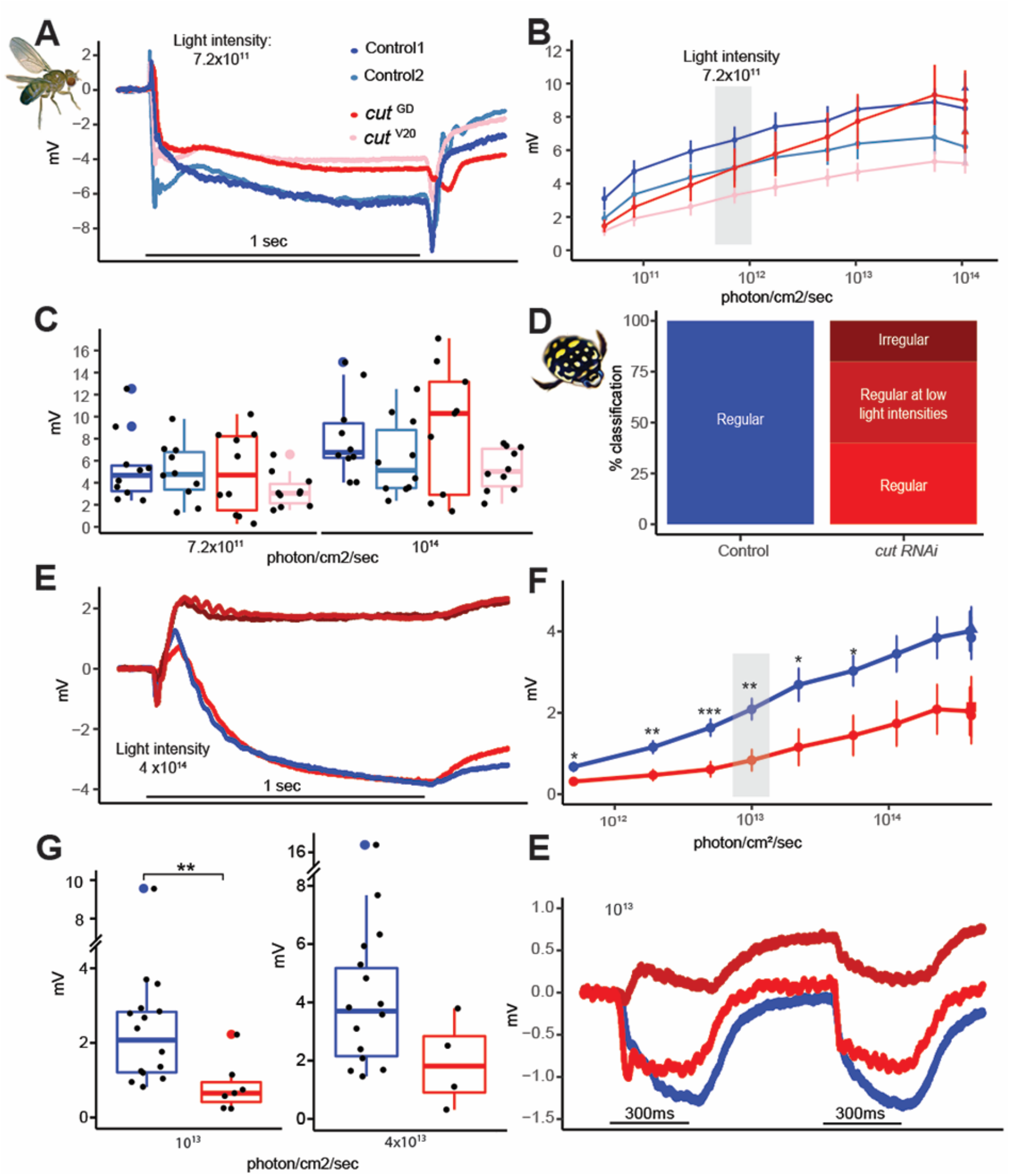
Despite major structural deficits, electroretinograms of *cut* knockdown individuals show relatively intact physiological responses in *D. melanogaster* and relatively minor deficiencies in *T. marmoratus*. A. Example recordings from two control and two test flies illustrate comparable responses. B. Average responses (with standard error) to increasing light intensities suggest a comparable dynamic range across the four tested fly lines (n = 10 each). C. Example responses at two different light intensities. D. Quantification of *cut*RNAi injected beetles shows inverted responses at all (dark red) or higher (medium red) light intensities. E. Example recordings of a control individual and one of each of the three phenotypes in (D). F. Average responses (with standard error) to increasing light intensities suggest a comparable dynamic range between control and *cut*RNAi individuals, albeit with generally lower responses in the knockdowns (*p < 0.05, ** p < 0.005; based on Wilcoxon’s rank sum test). G. Example responses at two different light intensities. H. Example of *cut*RNAi individuals showing different response dynamics when multiple pulses are presented.

It has previously been demonstrated that certain genetic perturbations can affect the ability of flies to maintain a proper photoresponse throughout a long series of light pulses (Riazuddin *et al*., 2012). To test if such deficiencies exist in *cut* knockdowns, we exposed flies and beetles to a series of 150 light pulses (extended sequence) (Supp Fig. 3). To assess differences in the PR potentials at earlier and later time points, the responses were analyzed by grouping the data into 10-point bins. Since some early signal reduction is normal (due to adaptation), we limited our analysis to bins 3-15. No significant differences between these bins were observed in any of the fly lines (Supp Fig. 3A,B, p = 0.97, 0.97, 0.44, and 0.58, Wilcoxon’s test) or the controls and *cut*RNAi beetles (Supp Fig. 3C,D, p = 0.29 and 0.28, Wilcoxon’s test). Consistent with our analysis at different light intensities (Fig. 7G), *cut*RNAi beetles showed slightly lower responses than the controls. Apart from some inverted responses in *cut*RNAi injected beetles (Fig. 7E,H), our ERG analysis did not reveal any major differences between the controls and knockdowns in either compound eye type.

## Discussion

We employed a comparative approach to test how Cut in SCs contributes to the development of two different compound eye types in insects. Our data support that this generally deeply conserved homeodomain transcription factor (CUX in vertebrates) is part of a deeply conserved gene network that is essential for proper eye development. The observed parallels are especially exciting because they are consistent with the idea that a common developmental pattern underlies diverse eye organizations (Lavin *et al*., 2022), and adds to a list of already known relatively well established deeply conserved genes, such as those of the RDGN (Gehring, 2001; Kumar and Moses, 2001; Mishra and Sprecher, 2020).

### Importance of Cut in SCs for precisely patterned compound eyes

Our comparative study suggests that if *cut* is knocked down, the crystalline precision of the eye is disrupted relatively early in development, irrespective of the compound eye type. The key to this phenomenon likely lies in the central role of SCs for ommatidium development and of Cut for proper patterning of SCs.

During eye development, SCs undergo dramatic changes in organization (Cagan and Ready, 1989) and play important roles in recruiting later-developing PPCs (Nagaraj and Banerjee, 2007) and regulating the orientation of developing PRs. The specific patterning steps of SCs likely include a combination of forces driven by the actomyosin cytoskeleton, cell adhesion (Cadherins and Nephrins), endocytosis, and Notch signaling (Chan *et al*., 2017; Blackie *et al*., 2020, 2021; Charlton-Perkins *et al*., 2021). As Cut expression is detectable shortly after SC specification (Blochlinger *et al*., 1990; Canon and Banerjee, 2003; Charlton-Perkins *et al*., 2011), it is well positioned to contribute to the regulation of these patterning events. Consistent with this possibility, in both species, *cut* knockdowns lead to defects in cell placement, including the presence of triads or displaced tetrads in the SC layer (Supp Fig. 1B) and the lateral displacement of adjacent PRs (Fig. 2A). Interestingly, these defects are reminiscent of those observed in the developing *D. melanogaster* mutant pupal retina for Pax2, Hibris, Roughest, Mastermind (a Notch signaling inhibitor), and Wingless (Grillo-Hill and Wolff, 2009; Cordero and Cagan, 2010; Charlton-Perkins *et al*., 2011; Blackie *et al*., 2021). Although it is well established that Cut interacts with wingless and Notch signaling in the wing disc (Micchelli *et al*., 1997), evidence suggests that it does not interact with wingless in the eye disc (Cordero and Cagan, 2010). Cut’s interactions with other genes in the retina therefore remain subject of further studies.

### Importance of Cut in SCs for accurate lens development

In *cut* knockdowns, we observed severe abnormalities in lens formation, with many parallel deformities in the two species that suggest a conserved overall function. Some differences, such as defects on the outer lens surfaces, are likely related to eye-type-specific differences in lens organization. Specifically, *D. melanogaster* eyes are characterized by biconvex lenses, whereas *T. marmoratus* eyes are characterized by plano convex lenses (Fig. 1). These differences may have caused the fly eyes to show rough lens surfaces with prevalent holes (blueberry phenotype), whereas the beetle eyes maintained relatively smooth outer surfaces with only occasional dimples (Fig. 3). However, knockdowns of *cut* in both species exhibited indentations on the inner lens surfaces as well as a generally disorganized lens arrays containing misshaped and fused lenses. As the inner surface is closer to the location of lens secretion (a process mediated by SCs and all pigment cells (Cagan and Ready, 1989; Wang *et al*., 2012; Chaturvedi *et al*., 2014; Stahl *et al*., 2017*b*)), it is not surprising that inner-surface deficits are more consistent between the two eye types. Similar accessory-cell-based lens defects have already been observed in several *D. melanogaster* eye mutants (Higashijima *et al*., 1992; Charlton-Perkins *et al*., 2011; Wang *et al*., 2022). Of specific interest are the rough eye mutants of *prospero* and *pax2*, which specify SCs combinatorially (Charlton-Perkins *et al*., 2011), and *bar*, which is required for PPC function (Higashijima *et al*., 1992). Both these mutants and our knockdowns have irregularly shaped lens arrays, individual lenses with holes (including the blueberry phenotype), and flattened and fused lenses. These commonalities highlight the complexity of the process of proper lens formation, which involves many genes that act synergistically in multiple cell types.

Based on the severity of lens defects observed in both *cut*RNAi species, it is not surprising that we also noted major optical defects. In both species, lenses failed to form sharp images or to focus on the same plane, with associated variability in image magnification (Fig. 4). Such optical assessments are a powerful but underutilized method for characterizing the functional deficits of lenses, although morphometric modeling (Wang *et al*., 2022) has been implemented as an alternative assessment method. The observed optical deficits likely have dramatic implications on the spatial resolution of both compound eye types. It remains an open question whether *cut* knockdowns also affect the optics of the crystalline cone (CC), another important extracellular optical component of superposition eyes (Warrant and McIntyre, 1993; Meece *et al*., 2021). The CC, which re-inverts the image to allow for an upright image at the level of PRs, is likely also formed (or at least contributed to) by SCs (Nilsson, 1989). Unfortunately, unlike in other beetles, such as fireflies, in *T. marmoratus*, the CCs are not retained when isolating lenses and hence cannot be optically assessed.

### Importance of Cut in SCs for PR placement and rhabdomere morphology, with minimal effects on PR function

Although Cut affects SCs, *cut*RNAi individuals of both species also exhibit severe morphological defects in adjacent PR cells. On a gross morphological level, this includes laterally displaced rhabdoms that are already apparent in early development (Fig. 2) as well as longitudinally displaced PR nuclei, shortened rhabdoms, and rhabdoms that extend through the basement membrane (Fig. 5). In *D. melanogaster*, the latter two phenotypes are particularly severe. Notably in that species the orientation of rhabdomeres turns during development (Ready, 2002). It is possible that observed phenotypes relate to perturbations of that process, a possibility that warrants further investigation. These observed differences could be related to variations in the knockdown severity, but could also be related to organizational differences between the two eye types, with rhabdoms in beetles originating much deeper in the eye but terminating more distal to the basement membrane than those in flies. Parallel defects were also observed in cross-sections at the ultrastructural level, with rhabdoms of both species showing a variety of morphological defects including splitting, variable sizing, and lateral displacements (Fig. 6). In *D. melanogaster*, where the open rhabdom facilitates checking for the presence of all rhabdomeres, a few ommatidia appear to be missing some rhabdomeres. However, they could also have been displaced to a different plane; evaluation of this possibility requires future 3D TEM analysis. Our data adds to a growing body of evidence that suggests that an intact SC–PR interface is necessary for proper PR placement (during early pupation) and rhabdom elongation (during mid to late pupation (Longley and Ready, 1995), and that these developmental patterns may be conserved between different PR types. It is particularly noteworthy that SC-specific *pax2* RNAi flies have very similar patterning defects (Fu and Noll, 1997; Charlton-Perkins *et al*., 2017), as might be expected if these two genes act through a common pathway. In contrast, Blimp-1 knockdown in SCs also leads to eyes with shortened rhabdoms(Wang *et al*., 2022), although they do not cross the basement membrane. Interestingly, even defects in surrounding support cells, such as PPCs, have been reported to result in similar rhabdom deficiencies in flies, seen in Bar mutants (Higashijima *et al*., 1992). Overall, these findings indicate that the accurate patterning of SCs is necessary to allow PRs to be properly shaped and positioned. It has already been demonstrated that ineffective cell adhesion in PRs is a major contributor (Longley and Ready, 1995; Izaddoost *et al*., 2002), a topic that warrants further exploration.

In line with our interpretation that structural PR deficits in *cut* knockdown flies and beetles are secondary to disturbances of the entire ommatidial unit, we found that the physiological response of PRs was relatively intact, although there were notable phenotype differences between the two species. In *D. melanogaster*, the *cut*RNAi flies had normal responses despite abnormal morphology, similar to SC-specific *pax2* knockdown flies (Charlton-Perkins *et al*., 2017). The possible physiological phenotypes in flies are worth further investigation, as our study was hampered by a relatively large variation in responses, likely due to minor differences in eye color, and more subtle physiological deficits may only become apparent when PRs are substantially challenged (Riazuddin *et al*., 2012; Charlton-Perkins *et al*., 2017).

In contrast to the behavior in flies, the PR responses in *cut*RNAi beetles were significantly lower than those in the controls for some light intensities and were even inverted in many cases (Fig. 7). This seemingly reversed phenotype is reminiscent of that of *repo* (a general glial marker) mutant flies (Xiong *et al*., 1994). Since the RNAi in *T. marmoratus* was not cell type specific (in contrast to that in *D. melanogaster*), it is conceivable that the observed species-specific differences are related to *cut*-dependent roles in other cell types such as the subretinal glia (Bauke *et al*., 2015). Interestingly, the reversed response was sometimes transient, being present only in the first pulse (Fig. 7). As the ERG is a field potential measured from the surface of the eye (Belusic, 2011), this waveform could be related to defects in the resistance barrier and the complex flow of currents in the relatively tight spaces of the eye rather than an altered PR response.

### A glimpse into the complex genetic basis of eye diversification

The data presented here identify Cut as part of a support-cell-specific conserved gene network that is essential for proper eye development in different compound eye types, with largely parallel phenotypes in *cut* knockdowns in flies and beetles. Interestingly, several defects observed here parallel those in SC-*pax2* mutants, thus providing supportive functional evidence for a *pax2*-*cut* model in SCs that is essential for proper cell adhesion, lens secretion, and PR morphology (Fig. 8). Here, we studied these knockdown phenotypes for the first time within the framework of a direct comparison between eye types, laying the groundwork for further investigations into this model and how it might regulate the development of different compound eyes. Since diverse compound eyes, including those with different optics, are composed of ommatidia with highly conserved cellular compositions (Nilsson, 1989; Paulus, 1979), it is not surprising that the disruption of a central cell type, such as SCs, has dramatic effects on the entire complex ommatidial unit.

**Fig. 8.**
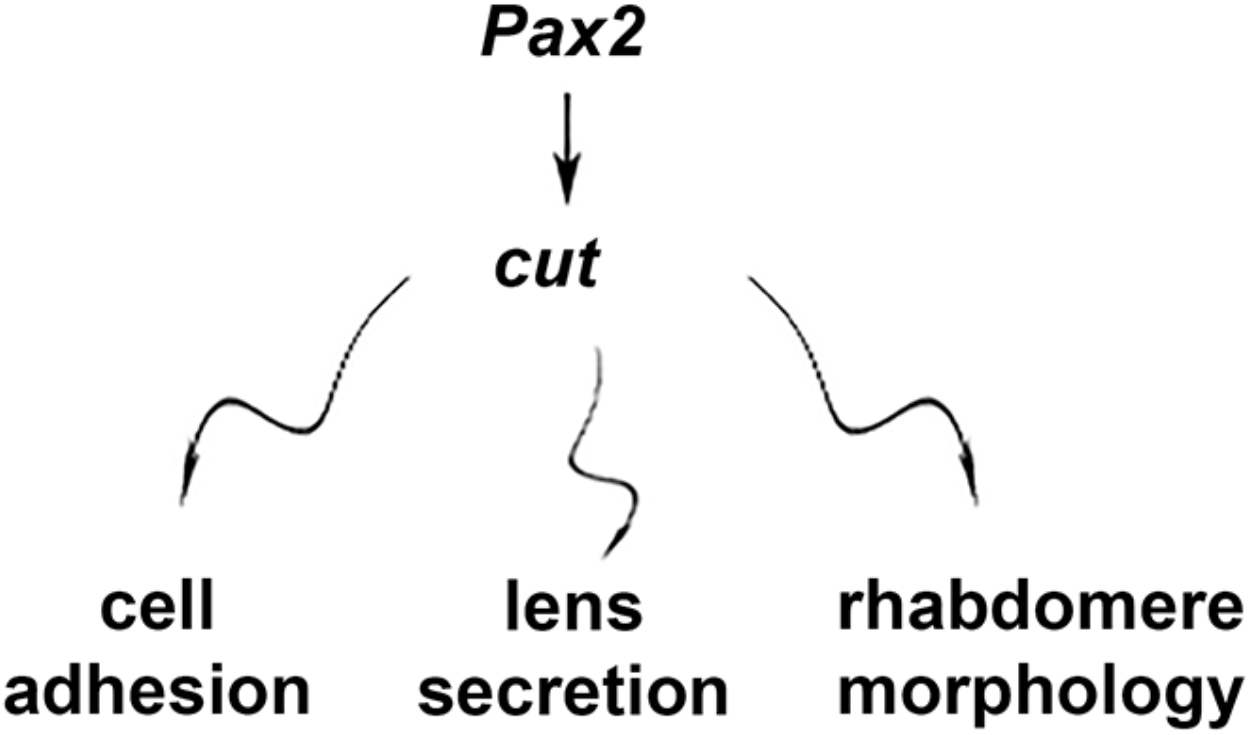
Schematic summary of SC-mediated effects of *cut*.

However, the manner in which related deficits manifest could be, at least to some degree, eye type specific. For example, we expect deficits to be particularly detrimental for the superposition optics of beetles, as a high level of precision in organization is required for proper function. Our data suggest that *cut*RNAi in beetles destabilizes this precision in multiple ways, including affecting ommatidial array regularity, lens integrity and optics, clear zone integrity, and PR placement. As each of these factors could itself be detrimental to proper function, we expect that even a relatively mild *cut* knockdown phenotype would lead to relatively dramatic visual deficits, with the potential of completely losing the ability to resolve images. For *D. melanogaster* eyes, the loss of spatial resolution is also expected, but this effect may be more subtle, as in parts of the eye, individual lenses would continue to project images (albeit possibly blurry images) on the corresponding PRs. Additionally, both eye types are typical for flying insects that capitalize on precise sampling from neighboring units for proper motion computation, which generally relies on elementary motion detectors (for a review, see (Egelhaaf *et al*., 1988)). Hence, even subtle phenotypes are expected to lead to specific deficits, which is an interesting area for further investigation.

Although this study compared two different optically relatively compound eye types, we have only scratched the surface of how Cut may be involved in the incredible diversity of eyes that have evolved in the lineage of compound eyes (Buschbeck, 2014; Morehouse *et al*., 2017; Lavin *et al*., 2022) and are composed of a set of highly conserved cell types (Paulus, 1979). One particularly intriguing observation in our study is the occurrence of fused lenses, which are particularly significant from an optical perspective, as such fusion events may give rise to eye formations in which one lens serves multiple PRs, the hallmark of a single-chamber, image-forming eye. Transitions from compound eyes to image-forming eyes have been observed in nature (for example, in mysid shrimp (Nilsson and Modlin, 1994)) but have thus far remained unexplored from a genetic perspective. We anticipate that our study is part of the groundwork for further explorations on support-cell-mediated eye diversification and how deeply conserved eye development genes beyond the RDGN contribute to the manifestation of optically highly diverse eye types.

## Materials and methods

### Animal husbandry and knockdowns

*D. melanogaster:* All flies were reared on standard cornmeal (made in-house) under a 12 h light–dark cycle at 27 °C in 60–70% humidity. Unless stated otherwise, adult flies were age controlled to 3 days old post-eclosion at the time of experimentation. For knockdown lines, the following alleles from the Bloomington Drosophila Stock Center (BDSC) and the Vienna Drosophila Resource Center (VDRC) were used: UAS-*cut*RNAi (GD4138)^VDRC1237^, UAS-*cut*RNAi (TRIP V20{HMS00924}attP2)^(BDSC 33967)^, UAS-*gfp*RNAi^(BDSC 9330)^, and UAS-mcherryRNAi (kindly provided by Dr. Vikki Weake, Purdue University (Stegeman *et al*., 2018). We used *pros*^*PSG*^*-GAL4 (Charlton-Perkins et al*., *2017)* to drive the UAS-*ct*^*RNAi*^ target in prospero-positive R7 and SCs. Flies with the following genotypes were used to generate *cut* knockdowns: 1) *yw*^67^; *pros*^PSG^-GAL4/*CyO*; UAS-*cut*RNAi/UAS-*cut*RNAi (*ct*^GD^), 2) *yw*^67^; *pros*^PSG^-GAL4/*pros*^PSG^-GAL4; UAS-*cut*RNAi/UAS-*cut*RNAi (*ct*^v20^), 3) *yw*^67^; *pros*^PSG^-GAL4/*pros*^PSG^-GAL4; UAS-*gfp*RNAi/UAS-*gfp*RNAi (control flies used in all experiments); and 4) *yw*^67^; *pros*^PSG^-GAL4/*pros*^PSG^-GAL4; UAS-mcherryRNAi/UAS-mcherryRNAi (a second control line for electrophysiology) (Fig. 7 and Supp Fig. 3).

*T. marmoratus:* The beetles used in this study were separated from our lab-grown colony at the third instar stage. Each larva was allowed to develop into an adult beetle in an individual pupation chamber filled with moist sand. All animals were reared under a 14 h light–10 h dark cycle at 25 °C. Unless stated otherwise, all adult beetles were ∼1 day old post-eclosion at the time of experimentation. To generate dRNAi against *cut*, we first identified the mRNA transcript of *cut* based on the three Cut domains and a homeobox domain (Supp Fig. 2 and Supp Table 1) from transcriptomics (Stahl *et al*., 2017*a*). To generate the probe, a 512 bp long unique region (outside the Cut and homeobox domains) was identified and amplified (see Supp Table 1 and Supp Fig. 2 for primers) from whole tissue cDNA. This amplicon was used for dsRNA synthesis as described in (Rathore *et al*., 2020). Then, 100 ng of *cut* dsRNA was injected in late-stage third instar larvae to ensure *cut* knockdown early during compound eye development (which is initiated during the pre-pupation stages in *T. marmoratus* larvae; personal observation).

### Immunohistochemistry

*D. melanogaster:* Larval, pupal, and adult eyes were processed as described (Charlton-Perkins *et al*., 2011) and stained with mCut (DHSB, 1:20), Elav (DHSB, 1:50), DCAD2 (E-cadherin; DHSB, 1:50), and drosocrystallin (kindly gifted by H. Matsumoto; rbCry 1:100; Fig. 1). The samples were imaged using a Nikon A1R multiphoton confocal microscope, and image processing was performed using NIS-Elements (Nikon) and Photoshop CC (Adobe). For Cut and N-cadherin staining, the pupal eye discs were dissected at ∼ 37% development as outlined in Tea et al. (Tea *et al*., 2014) In brief, tissue was fixed in 4% formaldehyde diluted in dissection solution for 20 min at RT. Post fixation, the eye discs were washed three times in PBT (PBS with 0.3% Tween 20). Then, 10% normal goat serum (NGS) in PBT was used to block the eye discs at RT. The eye discs were incubated in primary antibodies (anti-Cut 1:50 (DSHB) and anti-N-cadherin 1:50 (DSHB) in PBT) at 4 °C for ∼40 h. Anti-mouse Alexa Fluor 488 and anti-rabbit Alexa Fluor 488 were used as secondary antibodies at 1:500 for 2 h at RT. Subsequently, the eye discs were washed in PBT, the nuclei counterstained with DAPI, and after additional washing, mounted in Fluoromount (Fisher Scientific). Z-stacks were acquired using a Leica Stellaris 8 confocal microscope with a 40**×** objective at a resolution of 1024 **×** 1024 pixels and a pixel size of 0.047 μm.

*T. marmoratus*: Pupal eyes (at day 2 APF) were dissected and processed as described for *D. melanogaster* eyes, except that the tissue was counterstained with phalloidin (Alexa Fluor 647, Thermo Fisher) instead of anti-N-cadherin (due to a lack of cross-reactivity). Z-stacks were acquired using a Zeiss LSM 710, AxioObserver confocal microscope with a 40**×** objective at a resolution of 512 **×** 512 pixels and a pixel size of 0.11 μm. All images were processed using ImageJ, and the brightness and contrast were adjusted using Adobe Photoshop 2022.

### Cryosectioning and phalloidin staining

Control and *cut* knockdown fly and beetle heads were fixed in 4% formaldehyde overnight at 4 °C. These tissues were washed three times with PBS and then cryoprotected overnight at 4 °C in sucrose solutions of increasing concentrations (20%, 40%, and 60%). The samples were mounted in Neg50, flash frozen in liquid nitrogen, and cryosectioned at ∼18 μm (Leica CM1850). The sections were dried, rinsed in PBS, and stained with Phalloidin (following manufacturer’s instruction) and mounted with Fluoromount containing DAPI (Thermo Fisher). Z-stacks for *D. melanogaster* samples were obtained at a resolution of 1024 **×** 1024 pixels using a Leica Stellaris 8 confocal microscope with a 40**×** objective. *T. marmoratus* samples were imaged using a Zeiss LSM 710, AxioObserver confocal microscope with a 40**×** objective. The entire eye was tile scanned with a constant pixel size of 0.69 μm. All images were processed using ImageJ, and brightness and contrast were adjusted using Adobe Photoshop 2022.

### Differential interference contrast (DIC) microscopy and optical assessments

*D. melanogaster* and *T. marmoratus* heads for all groups were dissected in 100% and 50%, respectively, insect ringer (O’Shea and Adams, 1981) to maintain an appropriate osmotic environment and prevent the lens proteins from denaturing. Isolated lenses were suspended in corresponding insect ringer dilutions between coverslips to allow imaging of the undersurfaces of the lenses via DIC microscopy (Nomarski Optics, Olympus BX51 microscope with a 40**×** Uplan objective) as well as assessment of the optical quality of the lenses by visualizing the images produced by the lens array. The so-called “hanging drop method” follows protocols that were originally developed by (Homann, 1924) and since then commonly used in investigations of insect optics (Wilson, 1978; Buschbeck *et al*., 1999; Warrant *et al*., 2006; Stowasser *et al*., 2010). A representation of a 2 mm grating was projected through the lenses and pictures were taken of the resulting arrays of images that were focused by the lens arrays. All pictures were obtained using a microscope camera (Qimaging, Retiga 2000R) and the brightness and contrast were adjusted using Adobe Photoshop 2022.

### Electron microscopy (EM)

Transmission EM (TEM): *D. melanogaster* and *T. marmoratus* heads were dissected, fixed, and prepared for sectioning using standard protocols(Wolff, 2011) modified in the lab (Stowasser and Buschbeck, 2012). Images were acquired using a transmission electron microscope (JOEL JEM-1230 and Hitachi H-7650). The brightness and contrast were adjusted using Adobe Photoshop 2022.

Scanning EM (SEM): *D. melanogaster* and *T. marmoratus* heads were dissected, dried at -20 °C, and mounted on stubs with adhesive carbon pads (Electron Microscopy Sciences). The stubs were sputter coated with gold and imaged using a scanning electron microscope (FEI Apreo LV-SEM). The brightness and contrast were adjusted using Adobe Photoshop 2022.

### ERG measurements and statistical analysis

The genetic background in both groups resulted in increased variability in eye color. Since eye color influences the strength of ERG responses, the tested flies were visually color matched. As the two species have different spectral sensitivity peaks (Maksimovic *et al*., 2011; Sharkey *et al*., 2020), testing was carried out using a 490 nm light source for *D. melanogaster* and a 525 nm light source for *T. marmoratus*. The first test was used to assess the range of light intensities that led to PR responses (V–logI curves). This test consisted of a series of three 1 s long light flashes of increasing intensities, with each flash being ∼10 s apart to allow PR recovery. The second test was used to identify whether the PRs could sustain a response to a train of consecutive light flashes (extended sequence). This test consisted of a series of 150 light flashes at a single light intensity that was within the linear range of PR responses.

For *D. melanogaster* controls (*gfp*RNAi and *mcherry*RNAi) and *cut*RNAi (*ct*^GD^ and *ct*^V20^), flies were prepared and used for ERG measurements as previously described (Charlton-Perkins *et al*., 2017). A V–logI curve was established for each group using the following light intensities (photons/cm^2^/s): 4.28 **×** 10^10^, 8.15 **×** 10^10^, 2.78 **×** 10^11^, 7.20 **×** 10^11^, 1.75 **×** 10^12^, 5.42 **×** 10^12^, 1.04 **×** 10^13^, 5.53 **×** 10^13^, and 1.07 **×** 10^14^. To ensure steady recording, each series started with a pre-pulse of full light intensity. The extended sequence was administered at 7.20 **×** 10^11^ photons/cm^2^/s (the light intensity at which the PRs were ∼50% saturated), with light and interval durations of 300 ms. For *T. marmoratus* controls and *cut*RNAi, beetles were anesthetized using carbon dioxide. These beetles were secured on a glass slide with dental wax and then allowed to dark adapt for 10 min before establishing V–logI curves using the following light intensities (photons/cm^2^/s): 5.00 **×** 10^11^, 1.90 **×** 10^12^, 5.00 **×** 10^12^, 1.00 **×** 10^13^, 2.20 **×** 10^13^, 5.50 **×** 10^13^, 1.14 **×** 10^14^, 2.25 **×** 10^14^, 3.90 **×** 10^14^, and 1.07 **×** 10^14^. The recording stability was verified through a high intensity pre-pulse. The beetles were then allowed to dark adapt for 10 min before the extended sequence test was performed at 10^13^ photons/cm^2^/s (the light intensity at which the PRs were ∼50% saturated). As for flies, each light flash had a duration of 300 ms, however stimulus intervals were increased to 500ms to allow adequate PR recovery.

All the data were analyzed with a custom-made MATLAB code (Riazuddin *et al*., 2012; Charlton-Perkins *et al*., 2017). As some groups did not show a normal distribution according to the Shapiro-Wilks test, we used Wilcoxon’s rank sum test for all intergroup comparisons (in both species). All graphs were plotted and statistical tests were calculated in R (version 4.0.3, packages Dplyr, tidyr, and ggplot2).

## Acknowledgements

We thank Dr. John Mast for initial characterization of the *D. melanogaster ct*^*GD*^ phenotype and Dr. Aaron Stahl for the initial efforts in identifying and cloning *cut* in *T. marmoratus*; the beetle care team of the Buschbeck laboratory for helping with insect rearing; the Bloomington Drosophila Stock Center and Vienna Drosophila Resource Center for providing flies; Chet Closson and the Live Microscopy Core, College of Medicine (University of Cincinnati) for help with confocal imaging; the CCHMC Pathology Core for TEM imaging, with special thanks to Birgit Ehmer and Jessica Webster; Melodie Fickenscher and CEAC (University of Cincinnati) for SEM imaging; and Tamara Pace for editorial help. This research was funded by the National Science Foundation under IOS-1856241 (EKB), the National Institutes of Health grants R21-EY031526, EY024404, and unrestricted funds from Research to Prevent Blindness (TAC), and Cincinnati Children’s Hospital Medical Center (MCP).

## Author contributions

Conceptualization: S.R., E.K.B., T.A.C.; Investigation: S.R., M.C.P., M.M.; Data curation: S.R., E.K.B., T.A.C.; Writing - original draft: S.R., E.K.B.; Writing - review & editing S.R., E.K.B., T.A.C., M.C-P.; Supervision: E.K.B., T.A.C.; Funding acquisition: E.K.B., T.A.C.

## References

Agi E, Langen M, Altschuler SJ, Wu LF, Zimmermann T, Hiesinger PR. 2014. The evolution and development of neural superposition. Journal of Neurogenetics 28, 216–232.

Bauke A-C, Sasse S, Matzat T, Klämbt C. 2015. A transcriptional network controlling glial development in the Drosophila visual system. Development 142, 2184–2193.

Belušič, G. (2011). ERG in Drosophila. In Electroretinograms, (ed. G. Belušič): IntecOpen DOI:10.5772/21747.

Blackie L, Tozluoglu M, Trylinski M, Walther RF, Schweisguth F, Mao Y, Pichaud F. 2021. A combination of Notch signaling, preferential adhesion and endocytosis induces a slow mode of cell intercalation in the Drosophila retina. Development 148.

Blackie L, Walther RF, Staddon MF, Banerjee S, Pichaud F. 2020. Cell-type-specific mechanical response and myosin dynamics during retinal lens development in. Molecular Biology of the Cell 31, 1355–1369.

Blochlinger K, Bodmer R, Jan LY, Jan YN. 1990. Patterns of expression of cut, a protein required for external sensory organ development in wild-type and cut mutant Drosophila embryos. Genes & Development 4, 1322–1331.

Blochlinger K, Jan LY, Jan YN. 1991. Transformation of sensory organ identity by ectopic expression of Cut in Drosophila. Genes & Development 5, 1124–1135.

Blochlinger K, Jan LY, Jan YN. 1993. Postembryonic patterns of expression of cut, a locus regulating sensory organ identity in Drosophila. Development 117, 441–450.

Buschbeck EK. 2014. Escaping compound eye ancestry: the evolution of single-chamber eyes in holometabolous larvae. The Journal of Experimental Biology 217, 2818–2824.

Buschbeck E, Ehmer B, Hoy R. 1999. Chunk Versus Point Sampling: Visual Imaging in a Small Insect. Science 286, 1178–1180.

Buschbeck EK, Friedrich M. 2008. Evolution of Insect Eyes: Tales of Ancient Heritage, Deconstruction, Reconstruction, Remodeling, and Recycling. Evolution: Education and Outreach 1, 448–462.

Cagan RL, Ready DF. 1989. The emergence of order in the Drosophila pupal retina. Developmental Biology 136, 346–362.

Canon J, Banerjee U. 2003. In vivo analysis of a developmental circuit for direct transcriptional activation and repression in the same cell by a Runx protein. Genes & Development 17, 838–843.

Chan EH, Chavadimane Shivakumar P, Clément R, Laugier E, Lenne P-F. 2017. Patterned cortical tension mediated by N-cadherin controls cell geometric order in the eye. eLife 6.

Charlton-Perkins M, Cook TA. 2010. Building a fly eye: terminal differentiation events of the retina, corneal lens, and pigmented epithelia. Current Topics in Developmental Biology 93, 129–173.

Charlton-Perkins MA, Friedrich M, Cook TA. 2021. Semper’s cells in the insect compound eye: Insights into ocular form and function. Developmental Biology 479, 126–138.

Charlton-Perkins M, Leigh Whitaker S, Fei Y, Xie B, Li-Kroeger D, Gebelein B, Cook T. 2011. Prospero and Pax2 combinatorially control neural cell fate decisions by modulating Ras- and Notch-dependent signaling. Neural Development 6.

Charlton-Perkins MA, Sendler ED, Buschbeck EK, Cook TA. 2017. Multifunctional glial support by Semper cells in the Drosophila retina. PLoS genetics 13, e1006782.

Chaturvedi R, Reddig K, Li H-S. 2014. Long-distance mechanism of neurotransmitter recycling mediated by glial network facilitates visual function in Drosophila. Proceedings of the National Academy of Sciences of the United States of America 111, 2812–2817.

Cordero JB, Cagan RL. 2010. Canonical wingless signaling regulates cone cell specification in the Drosophila retina. Developmental Dynamics 239, 875–884.

Corty MM, Tam J, Grueber WB. 2016. Dendritic diversification through transcription factor-mediated suppression of alternative morphologies. Development 143, 1351–1362.

Cronin TW, Johnsen S, Justin Marshall N, Warrant EJ. 2014. Visual Ecology.

Ebacher DJS, Todi SV, Eberl DF, Boekhoff-Falk GE. 2007. Cut mutant Drosophila auditory organs differentiate abnormally and degenerate. Fly 1, 86–94.

Egelhaaf M, Hausen K, Reichardt W, Wehrhahn C. 1988. Visual course control in flies relies on neuronal computation of object and background motion. Trends in Neurosciences 11, 351–358.

Friedrich M. 2003. Evolution of insect eye development: first insights from fruit fly, grasshopper and flour beetle. Integrative and Comparative Biology 43, 508–521.

Fu W, Noll M. 1997. The Pax2 homolog sparkling is required for development of cone and pigment cells in the Drosophila eye. Genes & Development 11, 2066–2078.

Gehring WJ. 2001. The genetic control of eye development and its implications for the evolution of the various eye-types. Zoology 104, 171–183.

Grillo-Hill BK, Wolff T. 2009. Dynamic cell shapes and contacts in the developing Drosophila retina are regulated by the Ig cell adhesion protein hibris. Developmental Dynamics 238, 2223–2234.

Hardiman KE, Brewster R, Khan SM, Deo M, Bodmer R. 2002. The bereft gene, a potential target of the neural selector gene cut, contributes to bristle morphogenesis. Genetics 161, 231–247.

Hayashi T, Carthew RW. 2004. Surface mechanics mediate pattern formation in the developing retina. Nature 431, 647–652.

Higashijima S, Kojima T, Michiue T, Ishimaru S, Emori Y, Saigo K. 1992. Dual Bar homeo box genes of Drosophila required in two photoreceptor cells, R1 and R6, and primary pigment cells for normal eye development. Genes & Development 6, 50–60.

Homann H. 1924. Zum Problem der Ocellenfunktion bei den Insekten. Zeitschrift für Vergleichende Physiologie 1, 541–578.

Hsiung F, Moses K. 2002. Retinal development in Drosophila: specifying the first neuron. Human Molecular Genetics 11, 1207–1214.

Iyer J, Wang Q, Le T, et al. 2016. Quantitative Assessment of Eye Phenotypes for Functional Genetic Studies Using Drosophila melanogaster. G3 6, 1427–1437.

Izaddoost S, Nam S-C, Bhat MA, Bellen HJ, Choi K-W. 2002. Drosophila Crumbs is a positional cue in photoreceptor adherens junctions and rhabdomeres. Nature 416, 178–183.

Klann M, Schacht MI, Benton MA, Stollewerk A. 2021. Functional analysis of sense organ specification in the Tribolium castaneum larva reveals divergent mechanisms in insects. BMC Biology 19, 22.

Kumar JP. 2001. Signalling pathways in Drosophila and vertebrate retinal development. Nature reviews. Genetics 2, 846–857.

Kumar JP. 2012. Building an ommatidium one cell at a time. Developmental Dynamics 241, 136–149.

Kumar JP, Moses K. 2001. Expression of evolutionarily conserved eye specification genes during Drosophila embryogenesis. Development Genes and Evolution 211, 406–414.

Land MF, Nilsson D-E. 2012. Animal Eyes.

Lavin R, Rathore S, Bauer B, Disalvo J, Mosley N, Shearer E, Elia Z, Cook TA, Buschbeck EK. 2022. EyeVolve, a modular PYTHON based model for simulating developmental eye type diversification. Frontiers in Cell and Developmental Biology 10, 964746.

Longley RL Jr, Ready DF. 1995. Integrins and the development of three-dimensional structure in the Drosophila compound eye. Developmental Biology 171, 415–433.

Maksimovic S, Layne JE, Buschbeck EK. 2011. Spectral sensitivity of the principal eyes of sunburst diving beetle, Thermonectus marmoratus (Coleoptera: Dytiscidae), larvae. The Journal of Experimental Biology 214, 3524–3531.

Meece M, Rathore S, Buschbeck EK. 2021. Stark trade-offs and elegant solutions in arthropod visual systems. The Journal of Experimental Biology 224.

Meyer-Rochow VB. 2015. Compound eyes of insects and crustaceans: Some examples that show there is still a lot of work left to be done. Insect Science 22, 461–481.

Micchelli CA, Rulifson EJ, Blair SS. 1997. The function and regulation of cut expression on the wing margin of Drosophila: Notch, Wingless and a dominant negative role for Delta and Serrate. Development 124, 1485–1495.

Mishra AK, Sprecher SG. 2020. Early Eye Development: Specification and Determination. Molecular Genetics of Axial Patterning, Growth and Disease in Drosophila Eye, 1–52.

Morehouse NI, Buschbeck EK, Zurek DB, Steck M, Porter ML. 2017. Molecular Evolution of Spider Vision: New Opportunities, Familiar Players. The Biological Bulletin 233, 21–38.

Morrison CA, Chen H, Cook T, Brown S, Treisman JE. 2018. Glass promotes the differentiation of neuronal and non-neuronal cell types in the Drosophila eye. PLoS genetics 14, e1007173.

Nagaraj R, Banerjee U. 2007. Combinatorial signaling in the specification of primary pigment cells in the Drosophila eye. Development 134, 825–831.

Nepveu A. 2001. Role of the multifunctional CDP/Cut/Cux homeodomain transcription factor in regulating differentiation, cell growth and development. Gene 270, 1–15.

Nilsson D-E. 1983. Evolutionary links between apposition and superposition optics in crustacean eyes. Nature 302, 818–821.

Nilsson D-E. 1989. Optics and Evolution of the Compound Eye. Facets of Vision. Springer Berlin Heidelberg, 30–73.

Nilsson D-E. 2021. The Diversity of Eyes and Vision. Annual Review of Vision Science 7, 19–41.

Nilsson DE, Modlin R. 1994. A mysid shrimp carrying a pair of binoculars. The Journal of Experimental Biology 189, 213–236.

O’Shea M, Adams ME. 1981. Pentapeptide (proctolin) associated with an identified neuron. Science 213, 567–569.

Querenet M, Goubard V, Chatelain G, Davoust N, Mollereau B. 2015. Spen is required for pigment cell survival during pupal development in Drosophila. Developmental Biology 402, 208–215.

Paulus, H. F. (1979). Eye structure and the monophyly of the Arthropoda. In Arthropod Phylogeny, (ed. A. P. Gupta), pp. 299–384. New York: Van Nostrad Reinhold.

Rathore S, Hassert J, Clark-Hachtel CM, Stahl A, Tomoyasu Y, Bushbeck EK. 2020. RNA Interference in Aquatic Beetles as a Powerful Tool for Manipulating Gene Expression at Specific Developmental Time Points. Journal of visualized experiments: JoVE.

Ready DF. 2002. Drosophila compound eye morphogenesis: blind mechanical engineers? Results and Problems in Cell Differentiation 37, 191–204.

Riazuddin S, Belyantseva IA, Giese APJ, et al. 2012. Alterations of the CIB2 calcium- and integrin-binding protein cause Usher syndrome type 1J and nonsyndromic deafness DFNB48. Nature Genetics 44, 1265–1271.

Richter S. 2002. The Tetraconata concept: hexapod-crustacean relationships and the phylogeny of Crustacea. Organisms Diversity & Evolution 2, 217–237.

Scholtz G, Staude A, Dunlop JA. 2019. Trilobite compound eyes with crystalline cones and rhabdoms show mandibulate affinities. Nature Communications 10, 2503.

Schwentner M, Richter S, Rogers DC, Giribet G. 2018. Tetraconatan phylogeny with special focus on Malacostraca and Branchiopoda: highlighting the strength of taxon-specific matrices in phylogenomics. Proceedings of the Royal Society B. 285.

Sharkey CR, Blanco J, Leibowitz MM, Pinto-Benito D, Wardill TJ. 2020. The spectral sensitivity of Drosophila photoreceptors. Scientific Reports 10, 18242.

Stahl AL, Baucom RS, Cook TA, Buschbeck EK. 2017a. A Complex Lens for a Complex Eye. Integrative and Comparative Biology 57, 1071–1081.

Stahl AL, Charlton-Perkins M, Buschbeck EK, Cook TA. 2017b. The cuticular nature of corneal lenses in Drosophila melanogaster. Development Genes and Evolution 227, 271–278.

Stegeman R, Hall H, Escobedo SE, Chang HC, Weake VM. 2018. Proper splicing contributes to visual function in the aging Drosophila eye. Aging Cell 17, e12817.

Stowasser A, Buschbeck EK. 2012. Electrophysiological evidence for polarization sensitivity in the camera-type eyes of the aquatic predacious insect larva Thermonectus marmoratus. The Journal of Experimental Biology 215, 3577–3586.

Stowasser A, Rapaport A, Layne JE, Morgan RC, Buschbeck EK. 2010. Biological bifocal lenses with image separation. Current Biology: CB 20, 1482–1486.

Tea JS, Cespedes A, Dawson D, Banerjee U, Call GB. 2014. Dissection and mounting of Drosophila pupal eye discs. Journal of Visualized Experiments: JoVE, e52315.

Waddington, C. H. and Perry, M. (1960). The Ultra-Structure of the Developing Eye of Drosophila. Proceedings of the Royal Society of London B 153, 155–178.

Wang H, Morrison CA, Ghosh N, Tea JS, Call GB, Treisman JE. 2022. The Blimp-1 transcription factor acts in non-neuronal cells to regulate terminal differentiation of the Drosophila eye. Development 149.

Wang X, Wang T, Ni JD, von Lintig J, Montell C. 2012. The Drosophila visual cycle and de novo chromophore synthesis depends on rdhB. The Journal of Neuroscience 32, 3485–3491.

Warrant EJ. 1999. Seeing better at night: life style, eye design and the optimum strategy of spatial and temporal summation. Vision Research 39, 1611–1630.

Warrant EJ, Kelber A, Wallén R, Wcislo WT. 2006. Ocellar optics in nocturnal and diurnal bees and wasps. Arthropod Structure & Development 35, 293–305.

Warrant EJ, McIntyre PD. 1993. Arthropod eye design and the physical limits to spatial resolving power. Progress in Neurobiology 40, 413–461.

Wilson M. 1978. The functional organisation of locust ocelli. Journal of Comparative Physiology A 124, 297–316.

Wolff T. 2011. Preparation of Drosophila eye specimens for transmission electron microscopy. Cold Spring Harbor Protocols 2011, 1386–1388.

Xiong WC, Okano H, Patel NH, Blendy JA, Montell C. 1994. repo encodes a glial-specific homeo domain protein required in the Drosophila nervous system. Genes & Development 8, 981–994.

Yang X, Weber M, Zarinkamar N, et al. 2009a. Probing the Drosophila retinal determination gene network in Tribolium (II): The Pax6 genes eyeless and twin of eyeless. Developmental Biology 333, 215–227.

Yang X, Zarinkamar N, Bao R, Friedrich M. 2009b. Probing the Drosophila retinal determination gene network in Tribolium (I): The early retinal genes dachshund, eyes absent and sine oculis. Developmental Biology 333, 202–214.

