## Supplemental Figures for "Probing the conserved roles of Cut in the development and function of optically different insect compound eyes"

**Supplementary figures**

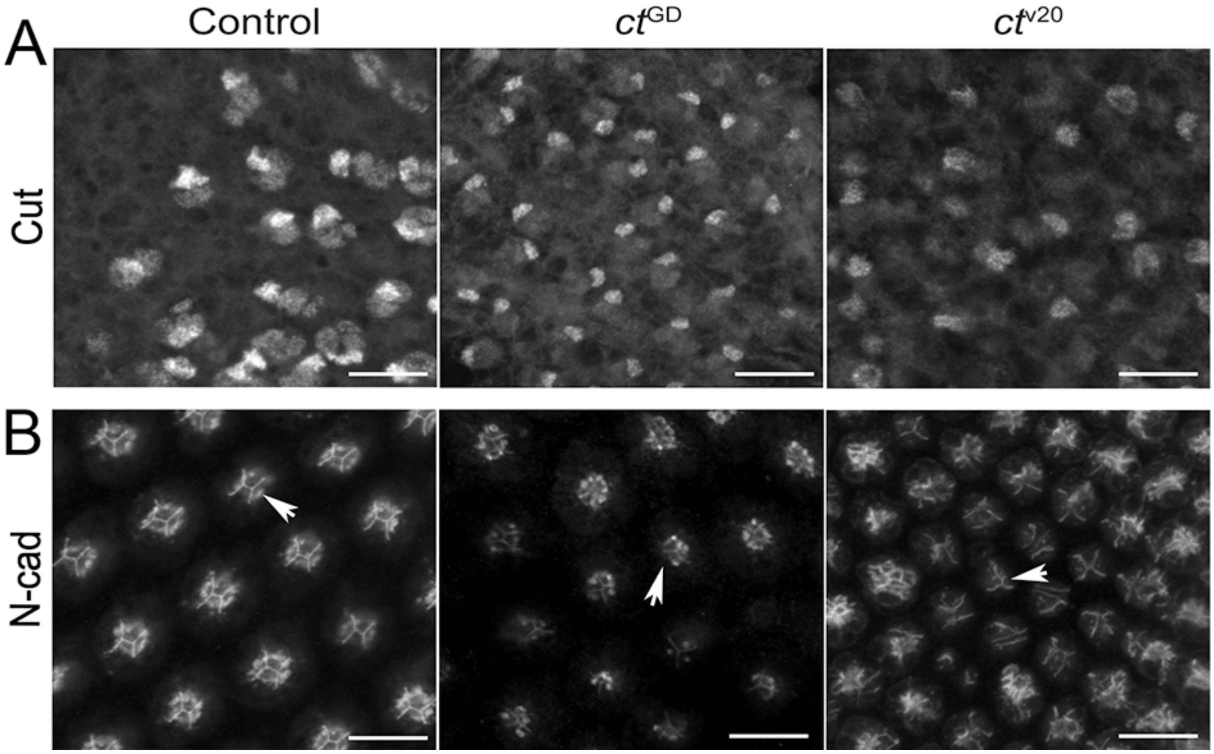


Supp Fig. 1. Confirmation of SC-specific *cut* knockdown in developing *D. melanogaster* eyes with knockdown-specific irregularities in the tetrad organization. A. To verify that *cut* knockdown was SC-specific, we also imaged Cut-positive sensory bristle nuclei and found comparable Cut immunoreactivity in all three fly lines. B. N-cadherin immunoreactivity illustrates that the control retinas show the typical “H” organization of the apical surfaces of the four SCs, whereas this organization is inconsistent or lost in both $ct^{GD}$ and $ct^{V20}$ retinas (arrows). Scale bars = 10 μm.

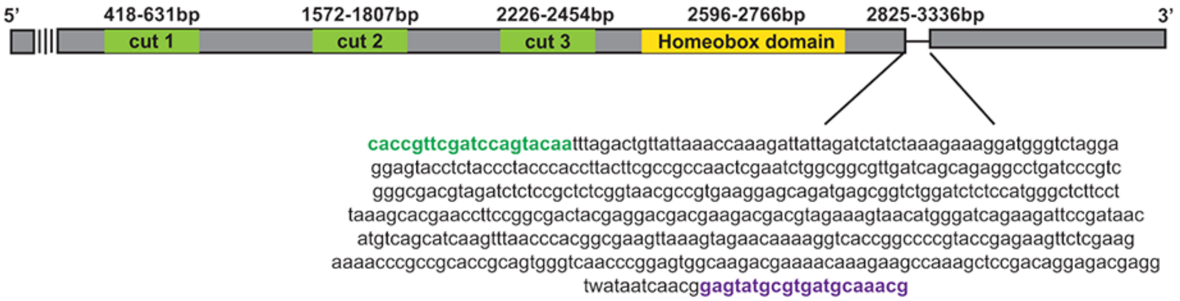


Supp Fig. 2. *Cut* gene in *T. marmoratus* and *cut*RNAi knockdown sequence. *Cut* is characterized by three specific domains (indicated as cut 1, 2, and 3) and a homeobox domain. The RNAi knockdown sequence is located toward the 3' end of the homeobox domain. The primer binding regions are indicated in green (forward) and purple (reverse).

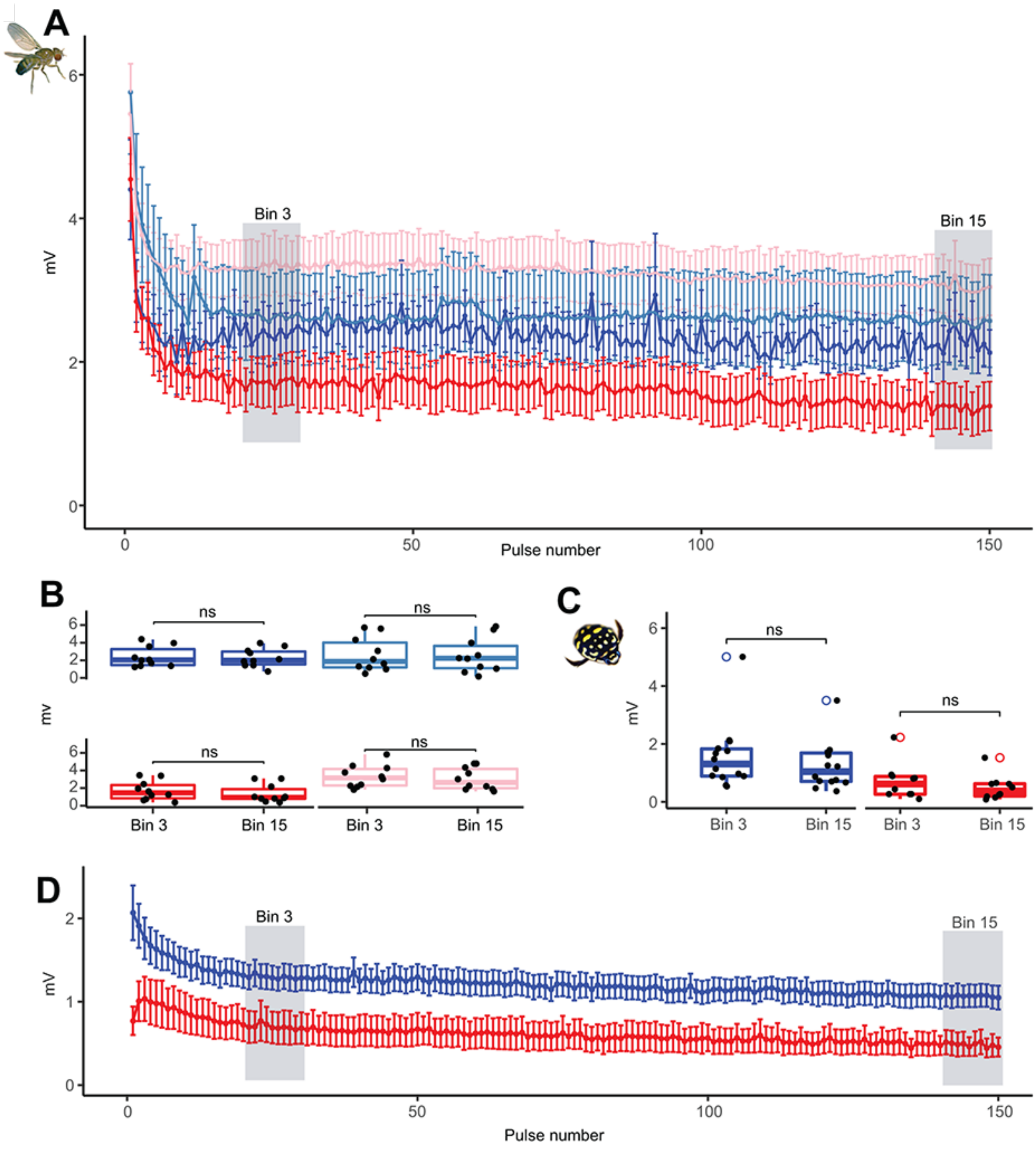


Supp Fig. 3. Electroretinograms for testing the sustainability of the photoreceptor response in both species. A. The average PR response (with standard error) of *D. melanogaster* to 150 pulses exhibits an initial decline (due to adaptation) followed by a somewhat lower signal than was sustained in all four lines throughout the pulse series. n = 10. B. Comparison between the responses of the third and last bins in *D. melanogaster*. C. Comparison between the responses of the third and last bins in *T. marmoratus*. D. PR response (with standard error) of *T. marmoratus* to 150 pulses, illustrating a sustained signal (after initial adaptation) in both control and *cut*RNAi individuals.

Supp Table 1. *cut* gene sequence identified from *T. marmoratus* transcriptome and the forward and reverse primer sequences used for amplifying the *cut* dsRNA amplicon.

| *cut* gene sequence 5’-3’ |
|---|
| GATGCTGTCCAAAAGATACCAACGCCAATACAAGCTGATTCTACCGAAAAATCTACTGCTTCTACCCCCGTACCTGAACAAA |
| CTGGAAGTCAGCCATCACCTCAACCTTCCCCGTTGATCAACGGGAATCCGAAGAGTCCAGAGGATAACAACAACGTACAC |
| GTGTCCAACAACAATATGAACCCACTCAACGCCGCAGCGATGTTATCCGTGAACAACTTCCTGAGGAACGACGATTCGCTG |
| AAGAGCCCTTACCGATTCGACGACCATAGATCACCGTTCAGATTCGCCGACGAACTCGGTATGCCTGGCGCAATGGTAGG |
| GAGACTAGGAGAAAGCTTAATCCCCAAAGGTGACCCAATGGAAGCAAGGCTGCAAGAGATGCTCCGCTATAACATGGATA |
| AATACGCAAACCAAAACCTGGACACGCTGACAATCTCCAGAAGGGTTCGCGAACTCCTGTCCATACATAACATCGGACAAA |
| GACTATTCGCTAAGTACATCCTGGGACTATCCCAGGGGACTGTTTCTGAGCTGCTGTCGAAGCCCAAGCCCTGGGATAAG |
| TTGACTGAGAAGGGACGGGACAGCTACAGGAAGATGCACGCATGGGCATGTGACGAGAATTCTGTCATGGTTCTCAAATC |
| GCTTATACCAAAAAAAGGTAAAGACACAACAATCCCCCCCTTCGGTCGACCCGAACCCGACATCACCGAGGAGAGGATCG |
| CCCACATCCTCAACGAAGCCACCAGTCACATGATGAAGTCGAACACTGAAGATTCGAGGAGCAACGATGACAGCAAAAGC |
| CCGTCTCAAGGACAGTGCCCAAGTCCCTTCTCCAAGGATTCTTCACAAAACAGAAGATTGAAGAAATACGAAAACGATGAT |
| ATACCTCAAGAAAAAGTAGTGAGAATATATCAAGAAGAATTAGCCAAGCTAATGGGAAGAAGAGTTGAAGATATGAGGAAT |
| CCAAGAGATCACTTTTCAAGTATCTTCTTCCCCCAATTCTTGAGCGGCGGTCTTCCAATGGACAGAACCCAAGAAGAAATC |
| CGGATGGCGTTAGACGCTTACCACAGAGAGCTGAGTAAGCTGAACCAGTGTCAGAATCCAGCTCAGCTTCCAAACCTCCC |
| AGGTCTTCCCGGTCTCCTGGCCCTCCAGCAGCAAGCTATGGTCCAGAACCCCCAACCTCAGAACGGCGGCGTCCAAGAC |
| TTATCCCTACCAAAAGACAACAAACCAAAGAGCATCAACGGCATGGAAGACGCGAAAGAAAAAGAAGCCATGGATGAAGC |
| CATGAAACACGCCGGCAGCGCGTTCAGCTTGGTCCGACCAAAACTAGAACCAGGAACCCAACAAGCAAGTTCCACAGCTT |
| CAAGTCCGCTAGGCAACTCCATTCTTCCACCTGCGATGACCCCAACAGATGACTTCACCTCATCAGCAGCCGCAAGCCCC |
| TTGCAGAGAATGGCTTCCATTACAAACTCACTGATCTCTCAACCACCGACCCAATCCCACCACAGTCCGGCACAGCGACCT |
| CTCAAAGCAGTTCTTCCTCCGATCACCCAACAACAGTTTGATCTCTACAACAACCTCAACACAGAAGACATCGTCAAGAAAG |
| TAAAAGAACAACTCAGTCAGTATTCCATCAGCCAGCGGTTATTCGGAGAAAGTGTCCTGGGTCTTTCGCAAGGATCAGTCA |
| GTGACCTCCTTGCGAGACCCAAGCCCTGGCACATGCTCACCCAGAAAGGACGCGAACCTTTCATCAGGATGAAGATGTTC |
| TTAGAAGATGAAAACGCAGTCCACAAACTGGTAGCCAGTCAGTACAAAATCGCACCCGAAAAACTCATGAGAACCGGCGG |
| ATACGGCACTTCCACCTCAGCTCTTGCCAAACCAATGCCACCCACGCCAAAAATGATCAGCGAAGCCGCCGGATTACTGA |
| ACAAAATGCAACAAGAAGGCCAAAATGCTCTTCTCCCTCCGAGCCTGAACCTTGGACCCCCTGGACAACAAAATCTTCCAC |
| AACCACCTCCCCCACCCATGTTGTTAACTCCTCCAGGAATCTCACCACATCATTCCATGTCCATGAAATCCGCCCAAGAAC |
| AAATGAAACAACAAAATCAAAGTCCTCATCCCGGAGTCCCCGTCTCACCGATGGGACAACAACCATCCTCCGCCATGAGAA |
| GTCTCCACCAACACATATCCCCAAGCGTCTACGAGATGGCAGCGCTAACACAAGACTTGGATACTCAAGTGATCACCACCA |
| AGATCAAAGAAGCTTTACTGGCCAACAATATTGGGCAAAAGATATTCGGGGAGGCCGTGCTGGGTCTTTCGCAGGGATCA |
| GTGAGCGAGTTGCTCTCAAAACCCAAACCTTGGCACATGTTGAGCATAAAGGGTCGCGAACCGTTCATCAGGATGCAACTT |
| TGGTTGAACGATGCTCATAACGTAGACCGCCTACAAGCACTGAAAAACGAAAGACGAGAAGCCAACAAAAGACGAAGATC |
| TTCTGGTCCAGGAGCTCACGATAACAGCTCGGACACGTCATCGAACGACACATCTGAGTTCTACCACTCCAACTCTCCAGG |
| ACCCGGGCCTCCATCCGCCAAGAAGCAGCGCGTGCTCTTCTCCGAGGAGCAGAAGGAAGCCCTCAGGTTGGCGTTCGCG |
| TTAGATCCGTACCCCAACGTGGCGACCATCGAGTTCCTGGCTGGCGAACTGGCCCTCAGCAGTAGGACCATCACCAACTG |
| GTTCCACAACCACCGTATGAGACTAAAGCAACAAGTCCCCCACGGAATGCCCTCCGACATCCCGCCGAGAGACCAAAACA |
| GCGGACAGACACCGTTCGATCCAGTACAATTTAGACTGTTATTAAACCAAAGATTATTAGATCTATCTAAAGAAAGGATGGG |
| TCTAGGAGGAGTACCTCTACCCTACCCACCTTACTTCGCCGCCAACTCGAATCTGGCGGCGTTGATCAGCAGAGGCCTGA |
| TCCCGTCGGGCGACGTAGATCTCTCCGCTCTCGGTAACGCCGTGAAGGAGCAGATGAGCGGTCTGGATCTCTCCATGGG |
| CTCTTCCTTAAAGCACGAACCTTCCGGCGACTACGAGGACGACGAAGACGACGTAGAAAGTAACATGGGATCAGAAGATT |
| CCGATAACATGTCAGCATCAAGTTTAACCCACGGCGAAGTTAAAGTAGAACAAAAGGTCACCGGCCCCGTACCGAGAAGT |

<table>
<tr><td colspan="2">TCTCGAAGAAAACCCGCCGCACCGCAGTGGGTCAACCCGGAGTGGCAAGACGAAAACAAAGAAGCCAAAGCTCCGACAG<br>GAGACGAGGTAATAATCAACGGAGTATGCGTGATGCAAACGGAAGACTACGGCAGACGGAATTCGGAAGAGACAGTTCG<br>GGTGGAACCTCAAGCGGTCATGGACCGGTTCGACGACGACGCGAGCGACGCGTCGTCGTCGGTGAGCCATGACGAGAA<br>CGCGGACCGTCGAAGCCCTTCTGCTCAAGTCAAACAAGAGAGAGAAGAACCTGAAGAAATAGTTACGAGACAAAGTAGTG<br>ATGAACAAATCGAACACACAGACAAACAAATTAAAACAGAGAACGAGGAAGAAAGATGGGAATATTAG</td></tr>
<tr><td>Forward: CACCGTTCGATCCAGTACAA</td><td>Reverse: CGTTTGCATCACGCATACTC</td></tr>
</table>
